# Optical TrkB activation in Parvalbumin interneurons regulates intrinsic states to orchestrate cortical plasticity

**DOI:** 10.1101/2020.04.27.063503

**Authors:** Frederike Winkel, Mathias B. Voigt, Giuliano Didio, Salomé Matéo, Elias Jetsonen, Maria Llach Pou, Anna Steinzeig, Maria Ryazantseva, Juliana Harkki, Jonas Englund, Stanislav Khirug, Claudio Rivera, Satu Palva, Tomi Taira, Sari E. Lauri, Juzoh Umemori, Eero Castrén

## Abstract

Activation state of Parvalbumin (PV) interneurons regulates neuronal plasticity, driving the closure of developmental critical periods and alternating between high and low plasticity states in response to experience in adulthood. We now show that PV plasticity states are regulated through the activation of TrkB neurotrophin receptors. Activation of an optically activatable TrkB (optoTrkB) specifically in PV interneurons switches adult cortical networks into a state of elevated plasticity within minutes by decreasing excitability of PV neurons. OptoTrkB activation induces changes in gene expression related to neuronal plasticity and excitability, and increases the phosphorylation of Kv3.1 channels. OptoTrkB activation shifted cortical networks towards a low PV configuration, promoting oscillatory synchrony and ocular dominance plasticity. Visual plasticity induced by fluoxetine was lost in mice lacking TrkB in PV neurons. Our data suggest a novel mechanism that dynamically regulates PV interneurons configuration state and orchestrates cortical networks during adulthood.

**Graphical Abstract:** 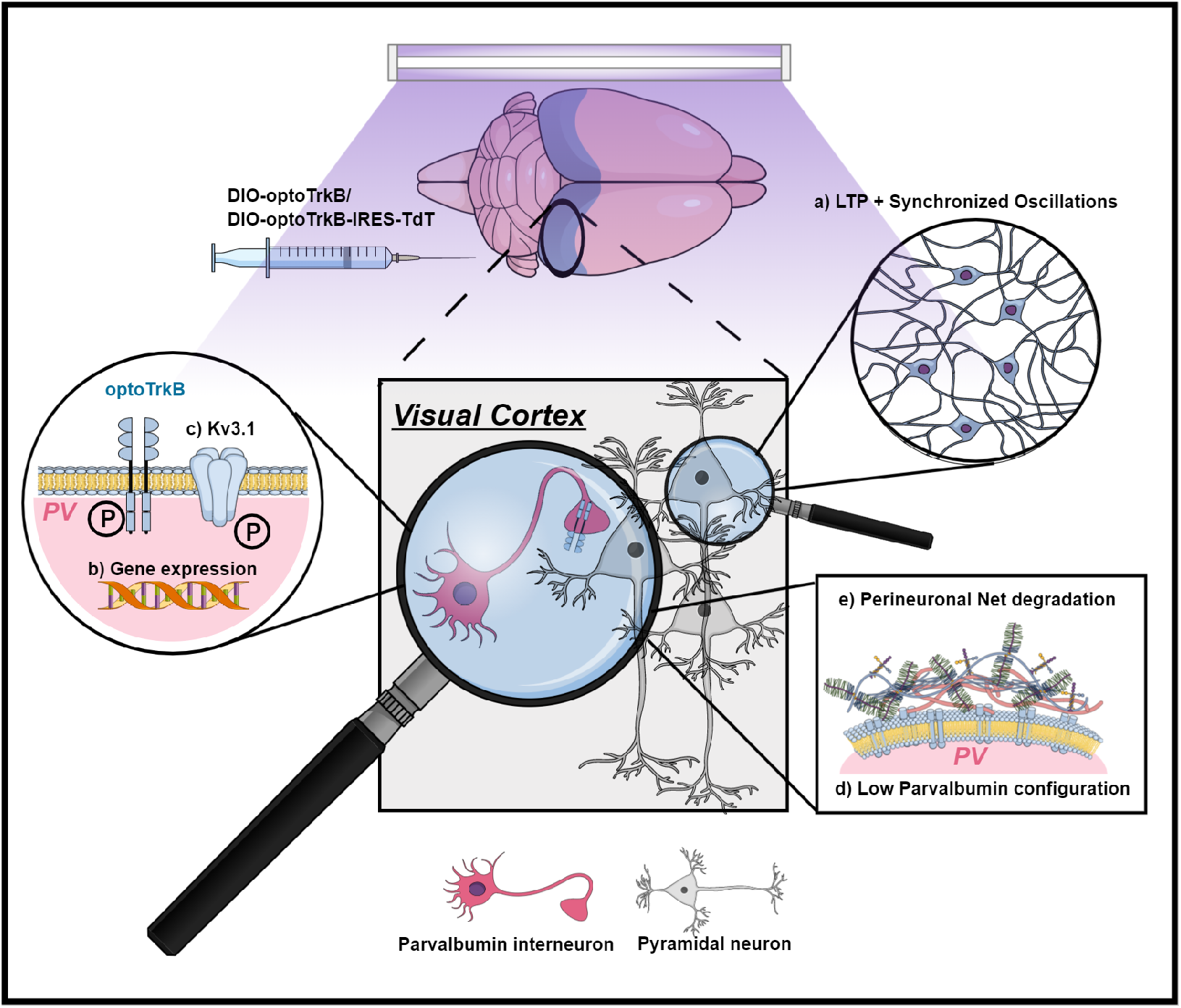

## Introduction

Brain plasticity is a key process to allow learning and adjust maladapted networks throughout life. The brain is particularly plastic during critical periods of early postnatal life (Hensch, 2005), and transition to a state of more limited plasticity in the adult brain coincides with the maturation of GABAergic parvalbumin-expressing (PV) interneurons. The role of the PV interneuron network has been best described for the critical period in the primary visual cortex, which is typically determined through ocular dominance plasticity (Huang et al., 1999; Jiang et al., 2010; Pizzorusso et al., 2002). PV interneuron maturation is promoted by brain-derived neurotrophic factor (BDNF) signalling (Huang et al., 1999) and the assembly of perineuronal nets (PNNs) (Pizzorusso et al., 2002), extracellular matrix components rich in chondroitin sulphate proteoglycans preferentially encasing PV interneurons. PV interneurons remain intrinsically plastic during adulthood, and external stimuli can switch the configurations of PV interneurons between plastic/immature and consolidated states as defined by low- and high-expression, respectively, of PV in these cells (Donato et al., 2013).

Increasing evidence has shown that a critical period-like state of plasticity can be evoked during adulthood by interventions, such as environmental enrichment (Sale et al., 2007) and antidepressant treatment (Karpova et al., 2011;Maya-Vetencourt et al., 2008; Mikics et al., 2018). Studies in the visual cortex have shown that adult ocular dominance plasticity is associated with decreased inhibitory activity thought to be driven by the PV interneuron network (Harauzov et al., 2010; Lensjø et al., 2017). However, the mechanisms underlying the switch in network states have remained a knowledge gap.

Considering that brain plasticity is an activity-dependent process that involves neurotrophin signaling, we hypothesized that the activity of the BDNF receptor neurotrophic receptor tyrosine kinase B (TrkB) within PV interneurons could regulate metabolic processes to mediate PV plasticity states during adulthood. TrkB is expressed in the majority of neurons, therefore, BDNF application to study TrkB functions in specific neuron populations holds considerable limitations (Gorba & Wahle, 1999; Kuczewski et al., 2009).

Here, we modified and optimized an optically activatable TrkB (optoTrkB) (Chang et al., 2014; Fenno et al., 2014) that can be expressed and activated in PV interneurons in a cre-dependent manner. This tool then allowed us to study the mechanisms underlying TrkB activity in PV interneurons and its effects on visual cortex plasticity as a well-defined model network. This optogenetic approach differs substantially from traditional channelrhodopsin experiments as the activation of optoTrkB allows us to control a relatively slowly developing, physiologically relevant molecular pathway.

We show here that optoTrkB activation specifically in PV interneurons in the primary visual cortex during 7 days of monocular deprivation (MD) was sufficient to reinstate ocular dominance plasticity. More strikingly, a single 30 seconds blue light stimulation of optoTrkB in PV interneurons resulted in LTP inducibility within 30 minutes and enhanced oscillatory synchrony in the visual cortex of adult mice. Subsequent single-nuclei sequencing revealed that activation of optoTrkB in PV interneurons rapidly induces changes in the expression of genes related to neuronal excitability and plasticity, such as *Grin1* and *Grik3*, subunits of the NMDA and kainate receptor, respectively. Surprisingly, also *PV* expression itself appeared to be regulated through TrkB activation. Consistently, we found a rapid decrease in PV intrinsic excitability and spontaneous excitatory postsynaptic current (sEPSC) frequencies after optoTrkB activation, as well as increased phosphorylation of the PV specific Kv3.1 channels. In line with our single-nuclei sequencing data, we found that TrkB activation in PV interneurons dynamically regulates PV and PNN configuration states, resetting the neuronal network into a plastic, immature-like state. Conversely, deleting TrkB from PV interneurons blocked these effects when induced pharmacologically using fluoxetine treatment. These findings demonstrate that TrkB activation in PV interneurons rapidly orchestrates cortical network plasticity by regulating intrinsic states and provide new evidence for its role in intracortical inhibition and plasticity modes during adulthood.

## Results

### TrkB activation in PV interneurons induces visual cortex plasticity

Using BDNF to study cell type-specific functions of TrkB has critical caveats as it can activate TrkB receptors in any neuron expressing it. To be able to specifically activate TrkB in PV interneurons, we optimized and modified an optically activatable TrkB (optoTrkB) where addition of a plant-derived photolyase homology region (PHR) domain into the C-terminus of full-length TrkB mediates light-induced dimerization of TrkB monomers (Chang et al., 2014) (Fig. 1A) (Figure S1A). We infected the V1 of adult mice specifically expressing cre recombinase in PV interneurons (PV-cre) with a lentivirus expressing a double-floxed inverted open reading frame of opto-TrkB (DIO-optoTrkB) to specifically express optoTrkB in PV interneurons (Figure S1B). Acute light activation of optoTrkB in V1 resulted in increased phosphorylation of TrkB and CREB (Figure S2A-S2E), suggesting successful activation of optoTrkB and downstream signalling *in vivo*.

**Fig. 1.**
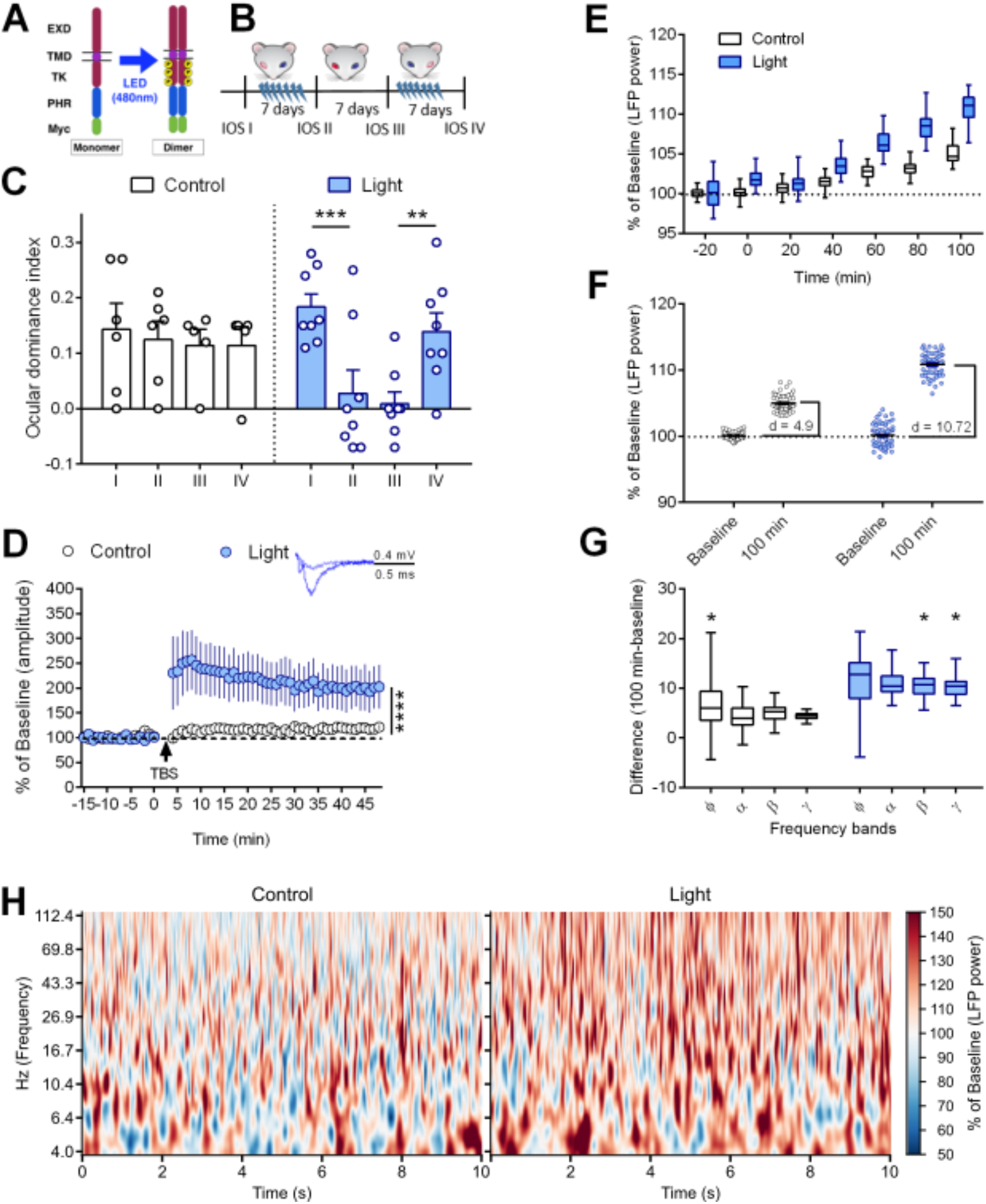
TrkB activation in PV interneurons induces visual cortex plasticity. (A) Structure of optoTrkB conjugated to a light reactive PHR domain, which dimerizes optoTrkB upon blue light exposure. B) Experimental timeline of the shift in ocular dominance paradigm with optoTrkB. OptoTrkB was stimulated with blue light twice daily (30 seconds) for 7 days during MD. (C) MD of the contralateral eye induces a shift in ocular dominance when combined with stimulation of optoTrkB (IOS I-II) (n = 6-8 animals/group). Two-way ANOVA with Sidak’s post-hoc test comparing the ODI of IOSI and II (control, p = 0.9094; light, p = 0.001). The shift is preserved if the eyes are not deprived (IOS II-III). Two-way ANOVA with Sidak’s post-hoc comparing the ODI of IOS II with III (control, p = 0.4985; light, p = 0.6915). Stimulation of optoTrkB reverses shift in ocular dominance when combined with MD of the ipsilateral eye (IOS III-IV) (n = 6-7 animals/group; one mouse excluded due to corneal infection). Two-way ANOVA with Sidak’s post-hoc test comparing ODI of IOS III and IV (control, p = 0.9094; light, p = 0.001). (D) LTP recordings from layer II/III in the V1. Theta-burst stimulation (TBS) results in LTP in slices where optoTrkB has been activated for 30 seconds 30 minutes prior to LTP induction, but not in control slices. Two-way ANOVA (p < 0.0001). n = 6-7 recordings/group randomly recorded from 5 animals. (E) The broadband (4–112 Hz) LFP power (blue: optoTrkB activated at 470 nm, black: control wavelength at 780 nm) averaged over animals (light = 5, control = 4) for each 20 minute recording session as a function of time after light stimulation. LFP power shows a significant increase over time (Regression analysis; Control: y = 0.04 [%/min]*x [min] + 100.25, R^2^ = 0.7573, p < 0.0001; Light: y = 0.09 [%/min]*x [min] + 101.01, R^2^ = 0.8240, p < 0.0001). (F) Broadband LFP power increase from baseline period (left plot of each half) to 100-minute after blue light (470nm) stimulation (right plot of each half) was approximately twice as strong after optoTrkB activation (blue, difference in medians: d = 10.72) than after stimulation with control wavelength (780nm) (black, difference in medians: d = 4.90). A Welch’s t-test confirmed that the LFP power in the 100-minute condition bins after optoTrkB activation is higher than after control stimulation (Welch’s t-test, t = 21.60, p < 0.0001). (G) Changes in frequency bands 100 min after light stimulation compared to baseline. (H) Wavelet spectra of control (left) and light stimulated (right) mice. Color scale. Flx, fluoxetine; ODI, ocular dominance index; MD, monocular deprivation; V1, primary visual cortex; PHR, photolyase homology region; EXD, extracellular domain; TMD, transmembrane domain; TK, tyrosine kinase domain. Bars represent means ± SEM. ** p < 0.01*** p < 0.001; **** p < 0.0001; Bars represent means ± SEM, **** p < 0.00001.

We then infected the V1 of adult PV-cre mice with DIO-optoTrkB lentivirus and subjected these mice to the standard protocol of ocular dominance (OD) plasticity induced by 7 days of monocular deprivation (MD) (see Material&Methods). As expected, mice infected with the lentivirus but having their transparent skull (Steinzeig et al., 2017) painted black to prevent light stimulation, failed to show any OD plasticity. In contrast, when the DIO-optoTrkB infected adult primary visual cortex (V1) was stimulated twice daily for 30 seconds by blue light through a transparent skull (Fig. 1B) during 7 days of MD, we observed a shift in ocular dominance index (ODI) towards the non-deprived ipsilateral eye (Fig.1C). The shift persisted for a week in the absence of visual deprivation or light stimulation (Fig. 1C). We then closed the eye that had been left open during the first MD session and again exposed the cortex to light twice daily for 7 days and again observed a shift towards the non-deprived eye (Fig. 1C). Light stimulation alone without DOI-optoTrkB virus infection during 7 days of MD had no effect on visual cortex plasticity (Figure. S2F).

A shift in ocular dominance takes several days to occur but we reasoned that direct activation of optoTrkB through light stimulation might have more immediate effects. We therefore studied the induction of long-term potentiation (LTP) as another proxy for visual cortex plasticity. LTP occurrence through theta-burst stimulation (TBS) in V1 is normally restricted to critical periods (Kirkwood et al., 1996). While TBS stimulation (Kirkwood & Bear, 1994) of acute V1 slices expressing optoTrkB in PV interneurons but kept in darkness did not induce LTP in slices, a robust LTP was induced in slices exposed for 30 seconds to blue light 30 minutes before the TBS stimulation (Fig.1D). These results suggest that direct activation of TrkB receptors in PV interneurons promotes excitatory transmission and plasticity within minutes.

Plasticity in the visual cortex is controlled locally by synchronized neuronal oscillations, which are regulated by the activity of PV interneurons (Galuske et al., 2019; Lee et al., 2012). *In vivo* local field potential (LFP) recordings of neuronal activity from V1 revealed a progressive increase in the broadband (4–112 Hz) LFP power in response to a 30-second long optoTrkB activation (Fig. 1E). The broadband magnitude at 100 minutes after stimulation showed a significant increase when compared to the pre-stimulation baseline. Furthermore, no increase in the broadband magnitude was observed in optoTrkB infected control animals stimulated with infrared light (780nm) (Fig. 1F, H). We also found increases in the magnitude at the separate frequency bands, particularly in alpha (α) and gamma (γ) range at 100 minutes after optoTrkB activation compared to baseline (Fig. 1G). Together, these results suggest that optoTrkB activation in PV interneurons promotes synchronized oscillations, which renders the network state of the visual cortex responsive and permissive for plasticity.

### TrkB regulates plasticity-related genes in PV interneurons

To identify the molecular mechanisms regulated by optoTrkB activation in PV interneurons, we performed single nuclei RNA sequencing (snRNA-seq) of visual cortical samples from DIO-optoTrkB-TdTomato (Figure S1C and S1D) transfected PV-cre mice 60 minutes after a 30-second light stimulation (Figure S3A). Single cell RNA sequencing was not feasible due to potential light-induced optoTrkB activation during cell extraction and processing (Figure S3B). Integrated data analysis (see Material&Methods) identified 18 clusters of neurons, including a cluster containing PV interneurons (Fig. 2A; Figure S4 and S5; Table S1-S3), which was further divided into four PV interneuron clusters using PCA analysis (Fig. 2B and 2C; Figure S6 and S7; Table S4). *Arc*, a known downstream signal of TrkB (Messaoudi et al., 2002, 2007; Yin et al., 2002; Ying et al., 2002), was only up-regulated in PV cluster 3 of the light-stimulated group, along with other immediate early genes, such as, *Jun* and *Egr1* (Fig. 2E), indicating that this cluster harbours optoTrkB transfected PV cells. In this cluster, genes related to dendrite morphogenesis, synaptic plasticity, and regulation of excitatory postsynaptic currents (EPSC) were differentially regulated in the light-exposed group (Fig. 2D) (Table S5). Strikingly, we found a decrease in *PV* expression itself among them (Fig. 2E), and this decrease was confirmed by qPCR (Figure S10). Particularly interesting is the downregulation of *Grin1* and *Grik3* that code for subunits of N-methyl-D-aspartate receptor (NMDA) and kainate receptors, respectively, and *Homeobox protein cut-like 2* (*Cux2*) (fold value, 0.63, Table S6), all being essential components of excitatory synapses (Fig. 2E). Furthermore, *Leucine Rich Repeat And Ig Domain Containing 2* (*Lingo-2*) that is broadly expressed in neurons was reduced after optoTrkB activation (fold value, 0.46, Table S6). Finally, we found an upregulation of the insulin-like growth factor 1 (IGF-1) receptor (*Igf1r*) expression (fold value, 1.29, Table S5), which shares many components with the BDNF pathway (Zheng & Quirion, 2004)

**Fig. 2.**
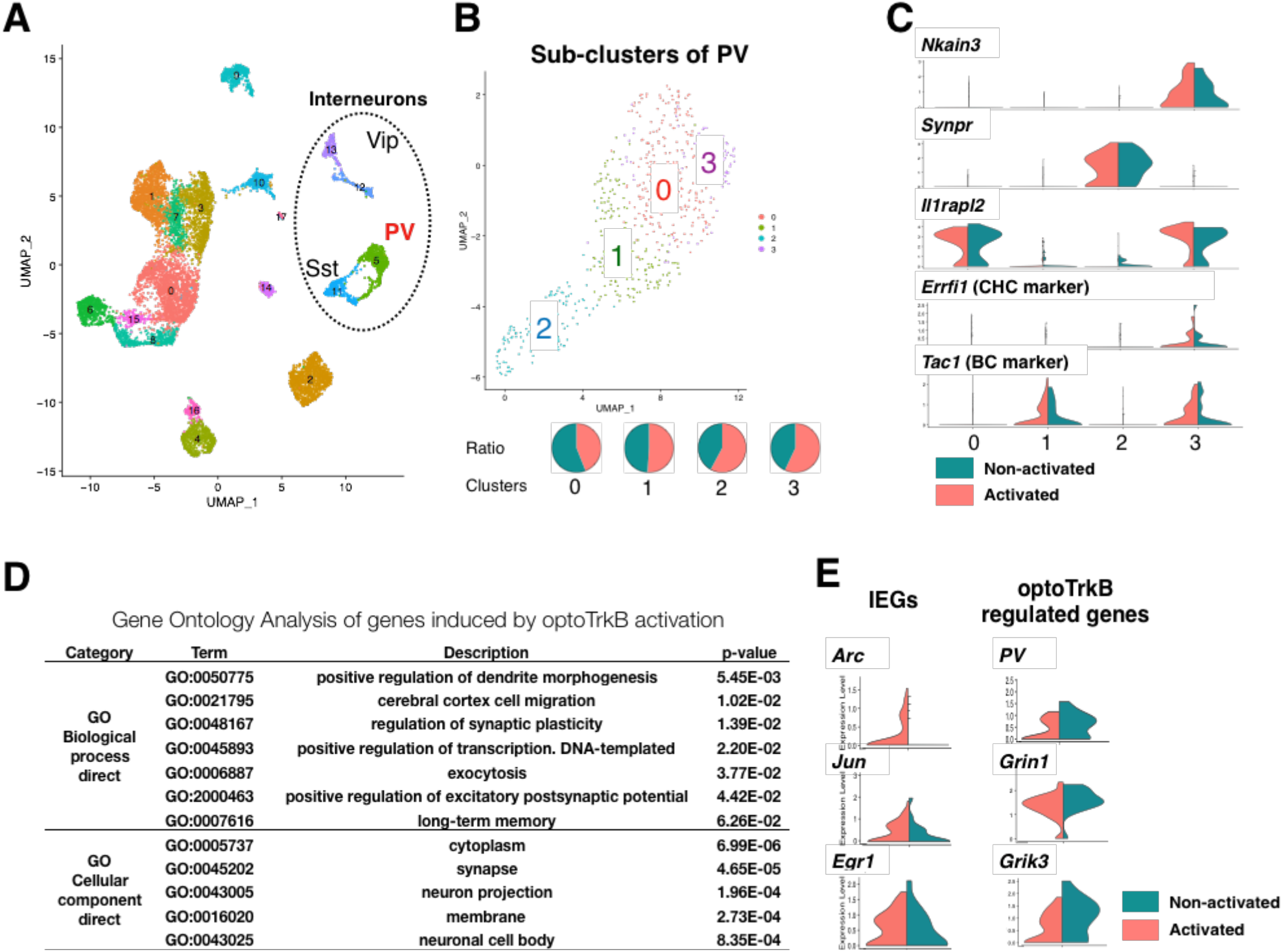
TrkB regulates plasticity-related genes in PV interneurons. (A) Two-dimensional Uniform Manifold Approximation and Projection (UMAP) plots based on 3,000 differentially expressed (DE) genes for 10,706 cells. After PCA analysis followed by clustering, the cells were grouped into 18 subpopulations including interneurons. (B) UMAP plots based on 3,000 DE genes for selected 586 cells clustered to PV interneurons. After PCA analysis, the cluster was further divided into 4 groups (PV_0, 1, 2, 3). (C) Markers differently expressed in each cluster. PV_3 cluster include both Chandelier (CHC) and basket (BS) cells. (D) Gene ontology (GO) analysis on differentially expressed (DE) genes between non-light-exposed and light-exposed samples in cluster 3_3. (E) Selected DE genes of immediate early genes (IEGs) and relevant genes to decreased excitation in cluster PV_3.

These results suggest that optoTrkB activation in PV interneurons changes the gene expression profile to particularly regulate the PV intrinsic properties.

### TrkB activation in PV interneurons reduces cell excitability

We next sought to validate the functional effects of optoTrkB activation on the intrinsic properties of PV interneurons. We therefore obtained patch clamp recordings from optoTrkB-positive PV interneurons in the V1 co-expressing TdTomato (Figure S1C and S1D). OptoTrkB was activated 10-60 minutes before the recordings using a 30-second blue light stimulation. We first recorded the intrinsic excitability by injecting current steps ranging from −100 to 600 pA. Strikingly, the intrinsic excitability of PV cells in optoTrkB stimulated slices was significantly lower 30-60 min after activation when compared to the non-activated controls (Fig. 3A and 3F) (Table S6A), and was accompanied by trends towards increased action potential (AP) half-width (Fig. 3B and 3F). We subsequently recorded spontaneous EPSCs (sEPSC) and found that the frequency of sEPSCs was also decreased in optoTrkB activated PV interneurons compared to controls (Fig. 3C and 3F), confirming functional changes in excitatory transmission onto PV interneurons. Neither the sEPSC amplitudes nor the frequency or amplitudes of spontaneous inhibitory postsynaptic currents (sIPSC) were changed (data not shown), suggesting that optoTrkB activation specifically affects excitatory inputs.

**Fig. 3.**
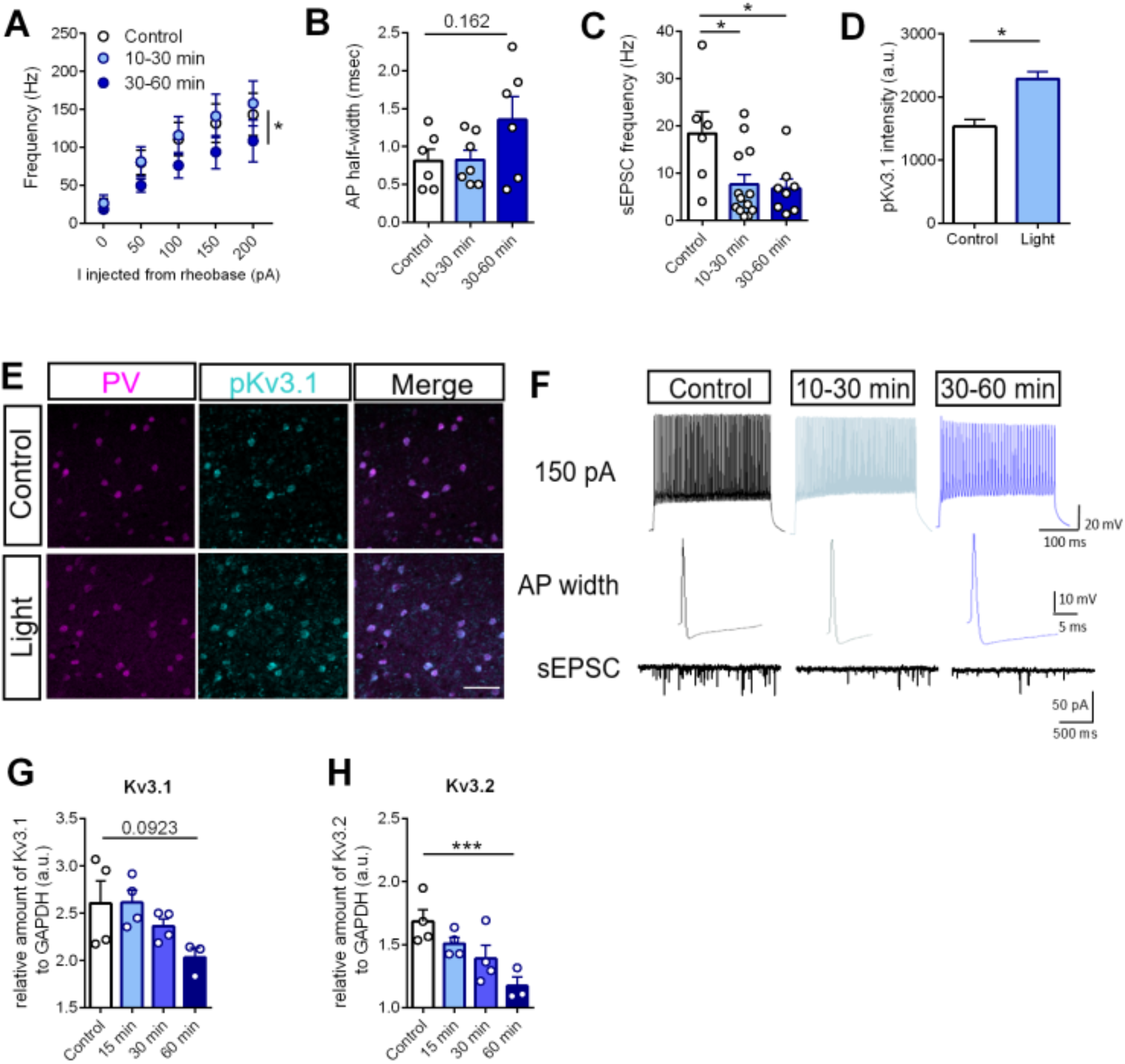
TrkB activation in PV interneurons reduces cell excitability. (A) Whole cell patch clamp recordings of intrinsic excitability of optoTrkB-transfected PV cells. Activation of optoTrkB decreases intrinsic excitability of PV cells at 30-60 minutes after light stimulation. Two-way ANOVA with Tukey’s post-hoc test comparing control with light stimulation (control vs. 30-60 min, p = 0.0354). n = 6-8 recordings/group randomly recorded from 14 animals. (B) Trends towards larger AP half-width 30-60 min after light stimulation. One-way ANOVA with Bonferroni’s post-hoc test (30-60 min, p = 0.1622). (C) sEPSC frequency in PV interneurons is lower at 10-30 minutes after light stimulation as compared to controls. One-way ANOVA with Bonferroni’s post-hoc test (10-30 min, p = 0.0233; 30-60 min, p = 0.0261). n = 6-8 recordings/group randomly recorded from 14 animals. (D) Light stimulation of optoTrkB results in increased intensity of phospho-Kv3.1 staining in PV interneurons in layer II/III of the V1. Unpaired t-test. n = 6 animals/group; 7 days twice daily light stimulation. (E) Representative images of PV, phospho-Kv3.1 and merged immunohistochemistry staining. (F) Representative traces used to estimate intrinsic excitability (top row, 150 pA), AP half-width (middle row) and sEPSC frequency (lower row). (G-H) qPCR quantification of Kv3.1 and Kv3.2 mRNA. Kv3.1 (p = 0.0923) (G) and Kv3.2 (H) mRNA levels are decreased 60 min after light stimulation measured by qPCR. qPCR measurements of control, 15 minutes, 30 minutes and 60 minutes after light stimulation. AP, action potential; spontaneous excitatory postsynaptic current, sEPSC; Bars represent means ± SEM. * p < 0.05; *** p < 0.0001; **** p < 0.00001

Potassium currents are known to regulate intrinsic excitability of neurons and Kv3 channels that are highly expressed in cortical PV interneurons regulate the fast-spiking properties (Chow et al., 1999; Du et al., 1996). Furthermore, Kv3.1 channels are directly inhibited by phosphorylation through PKC (Song & Kaczmarek, 2006), a downstream target of TrkB signaling. We therefore hypothesized that optoTrkB activation could result in enhanced phosphorylation or reduced expression of Kv3.1 channels. First, we immunohistochemically examined the expression intensities of phospho-Kv3.1 within PV interneurons and found that optoTrkB activation resulted in increased phospho-Kv3.1 expression in PV interneurons in visual cortical slices (Fig. 3D and 3E), which is consistent with reduced excitability. We then used qPCR with tissue samples from the V1 of optoTrkB-infected PV-cre mice 15 min, 30 min or 60 min after light stimulation to measure mRNA expression of Kv3.1 and Kv3.2 channels. The expression of Kv3.2 mRNA was significantly reduced and there was a progressive trend toward reduction for Kv3.1 mRNA (p = 0.0923) at 60 min after light stimulation (Fig. 3G and 3H). These data suggest that optoTrkB activation inhibits Kv3.1 potassium channels through phosphorylation and regulates Kv3.1 and Kv3.2 mRNA expression, thereby reducing the intrinsic excitability of PV interneurons.

### TrkB activation regulates PV-plasticity

PV expression itself is plastic and regulated by experience (Donato et al., 2013, 2015; Karunakaran et al., 2016), low PV expression being associated with plastic and high PV with consolidated networks. In addition, PV interneurons become gradually enwrapped by PNN during critical periods (Jiang et al., 2005) and PNN removal during adulthood reinstates a juvenile-like plasticity and reduces intracortical inhibition (Lensjø et al., 2017; Pizzorusso et al., 2002). As our snRNA-seq data revealed that optoTrkB activation decreases *PV* expression, we hypothesized that optoTrkB activation could directly mediate PV and PNN plasticity states. Indeed, activation of optoTrkB in the V1 PV interneurons for 30 seconds twice daily for 7 days reduced the PV intensities particularly in the subgroup of PV interneurons expressing high levels of PV (Fig. 4A, B) and the PNN intensities were also reduced in the same high-PV subgroup (Fig. 4C), suggesting a switch towards a low PV expressing network state. We next reasoned that a reduction in PNN intensities could correlate with a decrease in PNN expression and quantified the numbers of PNNs. We found that optoTrkB activation reduced the numbers of PNN-positive (PNN^+^), PV/PNN-double-positive (PV^+^PNN^+^) cells, and PV^+^PNN^+^ cells within the PV population (Fig. 4D-G), suggesting that PV interneurons switch to a critical period-like state after optoTrkB activation.

**Fig. 4.**
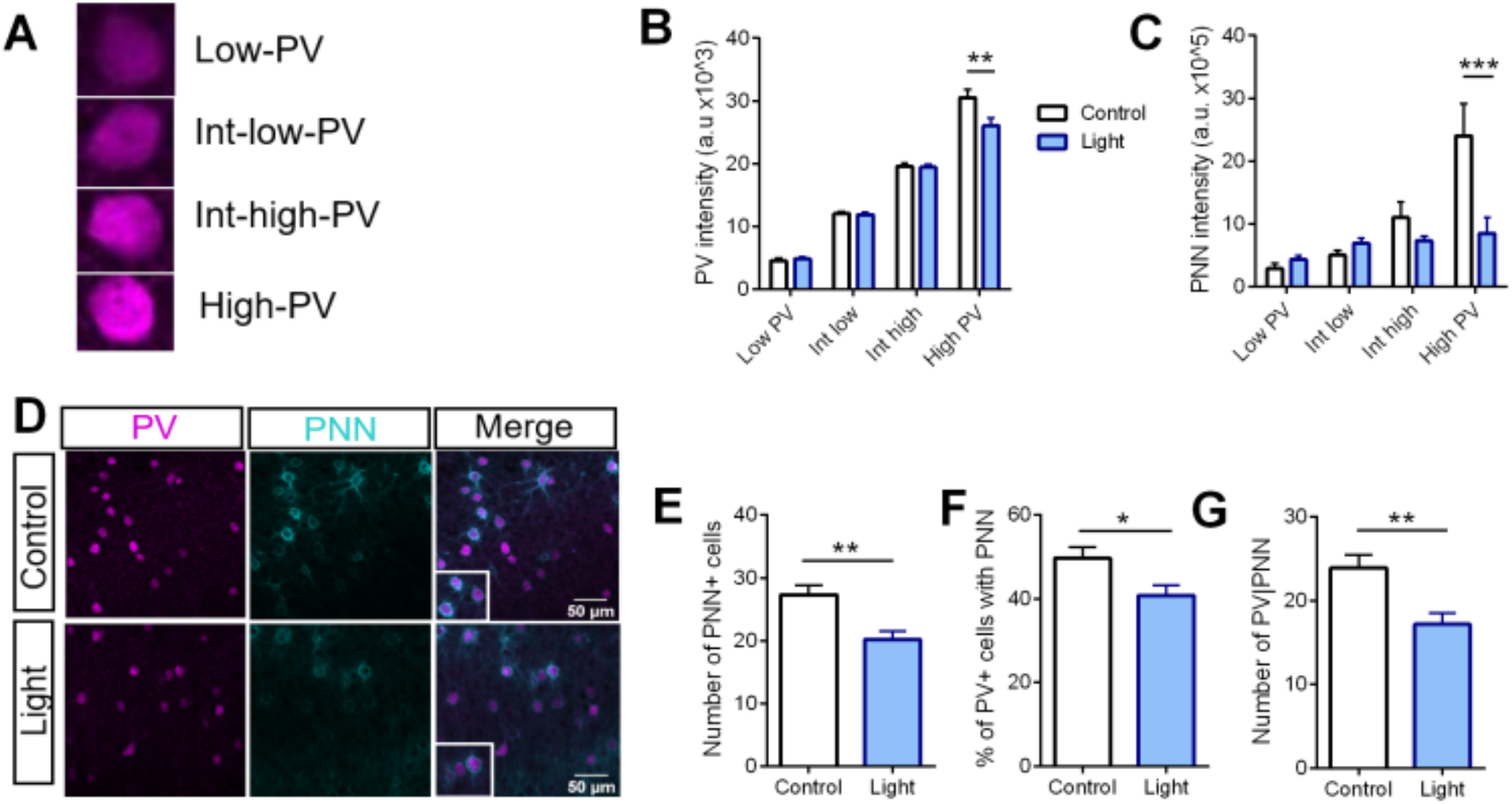
TrkB activation regulates PV-plasticity. (A) Representative immunohistochemical images of PV and PNN expression in layer II/III of the V1. (B-D) Number of cells expressing PV and/or PNN after optoTrkB activation (n = 6 animals/group). (B) Chronic light stimulation of optoTrkB significantly reduced numbers of PNN positive cells (Unpaired t-test; p 0.0012), (C) PV/PNN double-positive cells (Unpaired t-test; p = 0.0021), and (D) PV cells expressing PNN (Unpaired t-test; p = 0.0156). (E) Representative images of low, intermediate-low, intermediate-high and high PV-expressing cells, where PV interneurons are categorized according to PV intensity. (F) Image analysis on PV intensities. Chronic stimulation of optoTrkB significantly reduces the PV staining intensities of high PV expressing cells. Two-way ANOVA with Bonferroni’s post-hoc test comparing control vs. light stimulated samples (p = 0.0034). (G) Image analysis on PNN intensities. Activation of optoTrkB decreases PNN intensities in high PV-expressing cells. Two-way ANOVA with Bonferroni’s post-hoc test comparing control vs. light stimulated samples (p = 0.0003). Bars represent means ± SEM. * p < 0.05; *** p < 0.0001; **** p < 0.00001

### TrkB activation in PV interneurons is necessary for induction of visual cortex plasticity

Together, our findings demonstrate that TrkB activity in PV interneurons is sufficient to render the cortical network towards a plastic configuration state. We wondered, however, whether TrkB activity is also necessary for it. To test this hypothesis, we used a “loss-of-function” approach using conditional heterozygous mice with PV-specific TrkB knockout (hPV-TrkB CKO) and pharmacologically induced plasticity using chronic fluoxetine treatment. Using again the standard protocol for ocular dominance plasticity, we confirmed our previous findings that chronic fluoxetine treatment induced a shift of ocular dominance in combination with MD in the adult V1 of wild-type mice (WT) (Maya-Vetencourt et al., 2008; Steinzeig et al., 2017). However, no OD shift was seen in hPV-TrkB CKO mice in response to MD during chronic fluoxetine treatment (Fig. 5A). Consistently, chronic fluoxetine treatment also permitted LTP induction after TBS stimulation, as previously reported (Maya-Vetencourt et al., 2008), but this effect was absent in hPV-TrkB CKO mice (Fig. 5B). These data suggest that TrkB activation in the visual cortex is necessary for fluoxetine-induced adult plasticity.

**Fig. 5.**
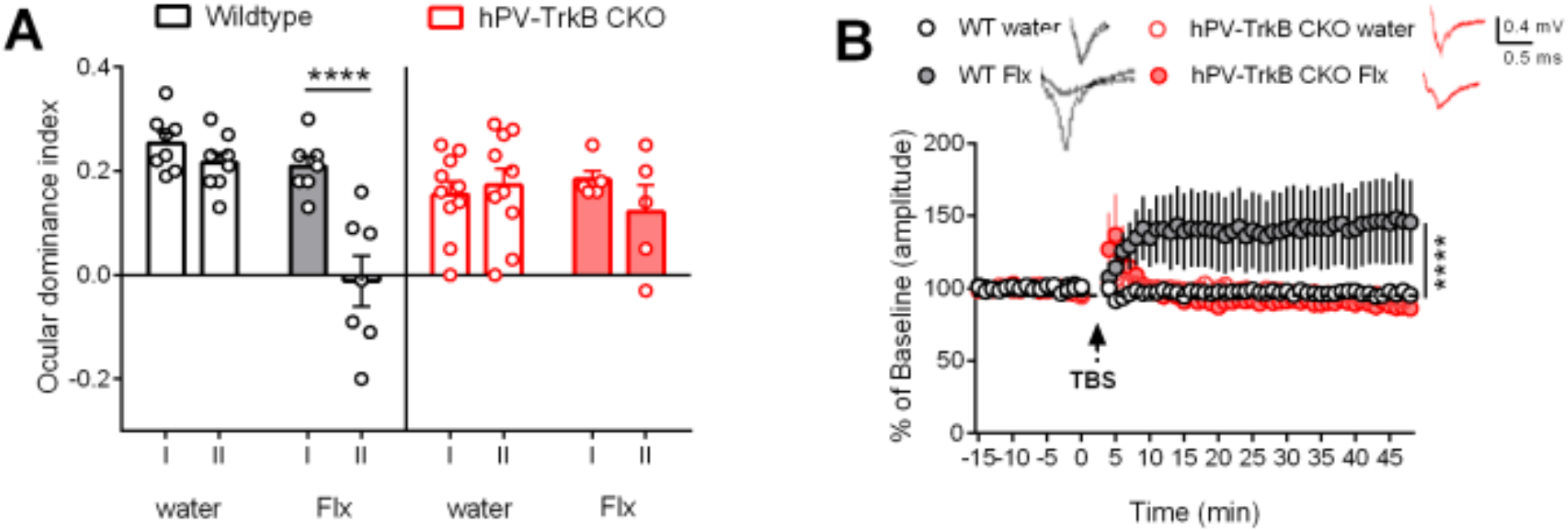
TrkB activation in PV interneurons is necessary for visual cortex plasticity. (A) ODI after chronic Flx treatment in WT and hPV-TrkB CKO mice. Flx permits MD to shift the ODI towards the non-deprived eye in the V1 of WT mice, but fails to do so in hPV-TrkB CKO mice. Two-way ANOVA with Sidak’s post-hoc test comparing the ODI of IOS I and II. WT water, p = 0.7311; WT Flx p < 0.0001; CKO water, p = 0.9621; CKO Flx, p = 0.5048. n = 5 – 10 animals/group. (B) LTP recordings from layer II/III in the V1 in WT and CKO mice after Flx treatment. TBS induces LTP only in WT mice treated with Flx. Two-way ANOVA with Tukey’s post-hoc test comparing treatment in WT and treatment in CKO mice (WT water vs. WT Flx, p = < 0.0001; CKO water vs. CKO Flx, p = 0.8871). n = 7-12 recordings/group from 3 animals.

### TrkB expression is necessary for fluoxetine-induced changes in PV intrinsic properties

We then sought to validate whether the underlying mechanisms of fluoxetine-induced plasticity are similar to those induced by optoTrkB activation. Indeed, similarly to what was observed after optoTrkB activation in PV interneurons, chronic fluoxetine treatment reduced intrinsic excitability (Fig. 6A, Fig.S7) (Table S6B), increased AP half-width (Fig. 6B, Fig.S7) and produced a trend towards a decrease in sEPSC frequency (p = 0.5397) (Fig. 6C, Fig.S7) in PV interneurons in WT mice, but none of these responses to fluoxetine were seen in hPV-TrkB CKO mice.

**Fig. 6.**
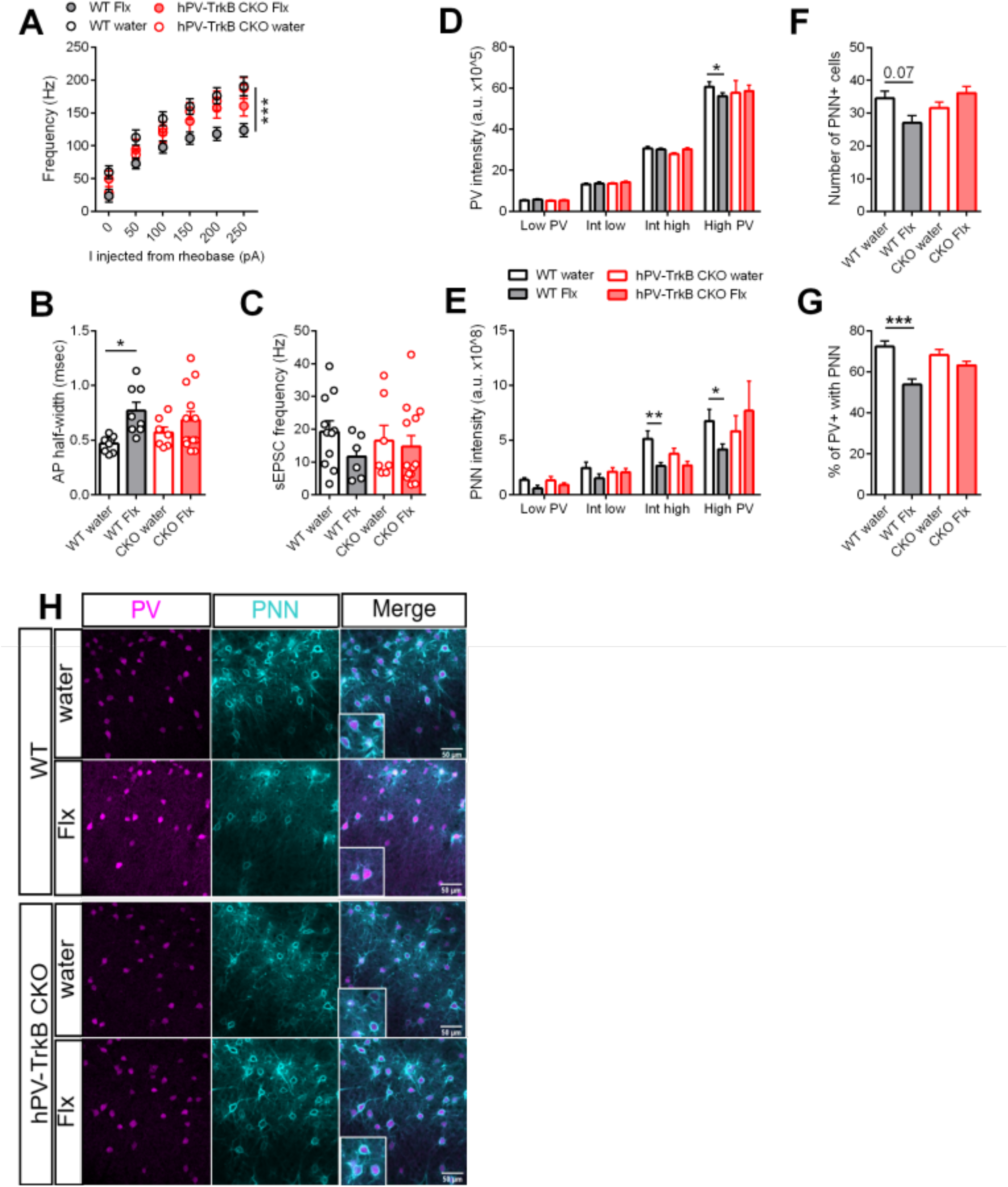
TrkB expression is necessary for fluoxetine-induced changes in PV intrinsic properties. (A) Whole cell patch clamp analysis of WT and hPV-TrkB CKO mice after chronic Flx treatment. The intrinsic excitability is reduced after Flx treatment in WT mice but not in hPV-TrkB CKO mice. Two-way ANOVA with Tukey’s post-hoc test comparing Flx treatment in WT and CKO mice (WT water vs. WT Flx, p < 0.0001; CKO water vs. CKO Flx, p = 0.7068). n = 7-12 recordings/group from 3 animals. (B) Flx treatment increases AP half-width in WT mice but not in hPV-TrkB CKO mice. Two-way ANOVA with Tukey’s post-hoc test comparing Flx treatment in WT and CKO mice (WT water vs. WT Flx, p = 0.03; CKO water vs. CKO Flx, p = 0.6672). (C) Flx treatment induces a trend towards reduced frequency of sEPSC in WT (p = 0.5397) but not in hPV-TrkB CKO mice (p = 0.9866). Two-way ANOVA with Tukey’s post-hoc test. (D) Flx treatment decreases PV intensities in high PV expressing cells in WT but this effect is not seen in hPV-TrkB CKO mice. Two-way ANOVA with Tukey’s post-hoc test comparing Flx treatment in WT and hPV-TrkB CKO mice (WT water vs. WT Flx, p = 0.0325; CKO water vs. CKO Flx, p = 0.9838). (E) Flx treatment reduces PNN intensities in intermediate-high and high PV expressing cells only in WT mice but fails to do so in hPV-TrkB CKO mice. Two-way ANOVA with Tukey’s post-hoc test comparing Flx treatment in WT and hPV-TrkB CKO mice (WT water vs. WT Flx, intermediate-high, p = 0.0014; high PV, p = 0.0325; CKO water vs. CKO Flx, intermediate-high, p = 0.3217; high PV, p = 0.6112). n = 6-10 animals/group. (F) Flx-treated WT mice show a trend towards lower numbers of PNN (p = 0.07). (G) Flx-treated WT mice have significantly lower percentages of PV interneurons also expressing PNNs, but this effect is abolished in hPV-TrkB CKO mice. Two-way ANOVA with Tukey’s post-hoc test comparing Flx treatment in WT and CKO mice (WT water vs. WT Flx, p = 0.0001; CKO water vs. CKO Flx, p = 0.6462). (H) Representative immunohistochemical images of PV and PNN expression in layer II/III of Flx-treated WT and hPV-TrkB CKO mice. Number (B, C) and intensity (D, E) of cells expressing PV and/or PNN by imaging analysis after Flx treatment. Bars represent means ± SEM. ** p < 0.01; *** p < 0.001; **** p < 0.0001

Finally, again in alignment with responses to optoTrkB, we found that chronic fluoxetine treatment reduced PV and PNN intensities among the high PV expressing population (Fig. 6D, E) and PNN^+^ abundance within the PV population in WT (Fig. 6F, G, H) but not in hPV-TrkB CKO mice (Fig. 6F-H). Taken together, these results strongly suggest that TrkB activation in PV interneurons is both necessary and sufficient to regulate PV intrinsic properties and switch the visual cortex to a state of high plasticity.

## Discussion

Our data show that TrkB signalling powerfully regulates the intrinsic activity of PV interneurons and thereby orchestrates cortical plasticity. TrkB activation in PV interneurons switches the intrinsic state of PV interneurons towards low excitability resulting in disinhibition and induction of critical period-like plasticity in cortical networks. Conversely, inhibition of TrkB signalling prevents induction of plasticity produced by fluoxetine treatment. Using optoTrkB as a methodological advancement we could show that TrkB particularly affects the expression of genes involved in synaptic transmission and intrinsic excitability as well as *PV* itself. The functional consequences of these changes are expressed in an increase of excitatory transmission, synchronized oscillations and visual cortex plasticity.

### TrkB activation in PV interneurons induces plasticity in adult brain

We previously demonstrated that chronic fluoxetine treatment can reactivate ocular dominance plasticity in the adult visual cortex (Maya-Vetencourt et al., 2008; Steinzeig et al., 2017). This effect was accompanied by a reduction in intracortical GABAergic transmission as measured by decreased extracellular basal GABA levels; induction of LTP after TBS stimulation and increased BDNF protein levels, strongly suggesting a regulatory role of intracortical inhibition in adult visual cortex plasticity (Maya-Vetencourt et al., 2008).

Although PV interneurons cover only a small percentage of the total neuronal population, their extensive axonal arborization enable strong inhibitory control over pyramidal cells (Hu et al., 2014). As BDNF promotes the maturation of PV cells during critical periods, we hypothesized that TrkB activation in PV interneurons could also affect plasticity processes during adulthood. Considering that administration of BDNF and fluoxetine activate all TrkB receptors expressed in the majority of neurons, we developed a cre-dependent optoTrkB to circumvent this problem and to be able to specifically activate TrkB signaling in PV interneurons only. Strikingly, while BDNF-induced TrkB activation during critical periods promotes the maturation of PV interneurons (Huang et al., 1999), TrkB activation in PV interneurons during adulthood reinstates a critical period-like plasticity machinery through rejuvenating these interneurons. These findings suggest that the plasticity machinery remains intact in the adult brain but needs the right tools to be reactivated.

### Differential effects of TrkB activation in pyramidal neurons and interneurons?

Reactivation of critical period-like plasticity in the adult brain is associated with reduced cortical inhibition and subsequent disinhibition of cortical networks (Hensch, 2005). BDNF has been shown to strongly promote neuronal activity, excitability (Figurov et al., 1995; Levine et al., 1995, 1998) and LTP (Kang & Schuman, 1994; Minichiello, 2009) in hippocampal and cortical excitatory neurons. In contrast, we found that optical activation of TrkB specifically in PV interneurons decreases excitability in PV cells and promotes field LTP in the visual cortex, indicating an increase in network-wide excitatory transmission and dynamics. These findings suggest that activation of TrkB produces differential effects on the cell excitability, increasing and decreasing it in pyramidal neurons and interneurons, respectively. Importantly, however, from the point of view of cortical networks, these differential effects on excitability cooperatively drive increased network activity, through enhanced excitability of pyramidal neurons and reduced activity of inhibitory neurons.

### Activation of TrkB mediates intrinsic changes in PV interneurons to enhance excitatory transmission in the visual cortex

The hallmark of PV interneurons is their high-frequency firing, which is enabled by the high expression of voltage-gated Kv3 channels (Chow et al., 1999; Hu et al., 2014). Kv3 channels are characterized by their fast deactivation during membrane repolarization, enabling sustained high-frequency firing. A decrease in Kv3 channel expression as well as their inhibition through PKC mediated phosphorylation is known to reduce cell excitability (Rudy et al., 1999; Song & Kaczmarek, 2006). PKC is a downstream target of TrkB signaling (Minichiello, 2009) and our data show that TrkB activation in PV interneurons results in enhanced phosphorylation of Kv3.1 and decreased expression of Kv3.1 and Kv3.2 channels, demonstrating control of TrkB signaling over PV intrinsic excitability. The decreased firing frequency of PV cells in turn enhances excitatory transmission necessary for visual cortex plasticity. The maturation of PV cells throughout critical period is concurrent with an increase in the fast-spiking firing frequency and an increase in Kv3.1 expression (Du et al., 1996; Plotkin et al., 2005). Interestingly, the fast-spiking properties of PV interneurons fails to develop in BDNF null mice (Itami et al., 2007). Considering that Kv3.1 channels are specifically expressed in high-frequency firing neurons, this could account for the differential effects of TrkB actions in pyramidal cells and PV interneurons.

### OptoTrkB-induced changes in gene expression in PV interneurons

Using snRNA-seq, we found genes that were differentially regulated after optoTrkB activation in PV interneurons. Particularly the expression of genes related to excitatory transmission, synaptic plasticity and excitability was affected. *Grin1* and *Grik3* are subunits of the NMDA and kainate receptor, respectively, and these essential components of excitatory synapses were reduced by optoTRKB activation. Knocking out *Grin1* in PV interneurons enhances network oscillations, particularly theta and gamma frequencies (Korotkova et al., 2010). The specific function of the *Grik3* subunit is not fully characterized, but kainate receptors have been shown to regulate both the maturation and excitability of GABAergic interneurons (Jack et al., 2019; Segerstråle et al., 2010), as well as the generation of synchronized activity at the level of neuronal networks (Bartos et al., 2007). As both NMDA and kainate receptors are known to mediate and modulate excitatory synaptic transmission (Zhang et al., 2013), the downregulation of *Grin1* and *Grik3* might therefore contribute to the changes in intrinsic properties of PV interneurons. In addition, we observed a decrease of *Cux2*, deletion of which reduces the amplitude and lowers the frequency of mEPSC (Cubelos et al., 2010). Furthermore, *Lingo-2* that is broadly expressed in neurons was reduced after optoTrkB activation, and the *Lingo* family members act as negative regulators of TrkB signalling (Meabon et al., 2016). Finally, we found an upregulation of the *Igf1r* expression. IGF-1 is another neurotrophic factor and shares components with the intracellular pathway of BDNF (Zheng & Quirion, 2004). Studies from the visual cortex have shown that IGF-1 reactivates ocular dominance plasticity and reduces GABAergic transmission in the adult visual cortex (Maya-Vetencourt et al., 2012). Upregulation of the IGF-1 receptor could render PV interneurons more responsive to IGF-1, thereby further contributing to a decrease in GABAergic transmission.

### Activation of optoTrkB regulates PV plasticity

Exciting research suggests that PV interneurons show intrinsic plasticity and can change between high and low plasticity states when exposed to different environmental conditions (Donato et al., 2013). For example, environmental enrichment that is known to promote neuronal plasticity, induces a low PV-expressing network state. In contrast, contextual fear conditioning that is associated with network consolidation, shifts the PV network into a high-PV expressing state (Donato et al., 2013). Additionally, PV expression progressively increases throughout development and could therefore also account for the reduction in brain plasticity during adulthood (Donato et al., 2013; Umemori et al., 2015). Interestingly, TrkB activation in PV interneurons can regulate PV plasticity states, and this effect may be mediated through direct regulation of PV mRNA expression as confirmed by single nuclei sequencing and qPCR. As PV is a Ca^2+^ buffer, a low PV expressing state could also directly regulate PV interneuron firing (Eggermann & Jonas, 2012; Hu et al., 2014)

In the hippocampus, PV intensity is generally higher in PV cells enwrapped by PNNs and PV cells with weak staining intensity often lack PNNs (Yamada et al., 2015). The maturation of the PV network is associated with the formation of PNNs (Pizzorusso et al., 2002). Enzymatic digestion of PNNs restores plasticity in adulthood (Lensjø et al., 2017; Pizzorusso et al., 2002) and results in a decrease in PV intensity and PV mRNA levels, suggesting a correlation between PNN expression and PV configuration states (Yamada et al., 2015). In fact, a recent study by Devienne et al. demonstrates that transient electrical silencing of visual cortical PV interneurons induces a regression of PNN (Devienne et al., 2019), indicating a causal relationship between PV activity and the accumulation of PNNs. Our data suggest that TrkB activation within PV interneurons directly regulates PV expression and leads to the reduction in PNN levels, thereby contributing to the plasticity state of PV neurons.

In conclusion, we show that TrkB receptor activation within PV interneurons is sufficient and necessary to rapidly reduce their excitability and plasticity state, thereby changing cortical network dynamics. Hence, TrkB activity in PV cells dynamically controls network rewiring, learning and consolidation. Although we used the visual cortex as a model network, these mechanisms might be extrapolated to other brain areas, such as the hippocampus and amygdala, and therefore aids in the understanding of the pathophysiology of neuropsychiatric disorders and the rational design of clinical interventions.

## Acknowledgments

We thank Drs. Beatriz Rico and Oscar Marín for their comments on the manuscript. We thank our lab technicians Sulo Kolehmainen and Outi Nikkilä for technical and practical help to realize the experiments. We also thank the animal caretakers in the Laboratory Animal Center of the University of Helsinki (UH) for help and support with the animals, and Louisa Böttcher, Mirko Torheiden and Marvin Krampe for their help with the Western Blots. We also thank Noora Aarnio and Nina M Peitsaro in the Biomedicum Flow cytometry unit for helping FACS, and Jenni Lahtela and Bshwa Ghimire in FIMM Single Cell Analytics core facility supported by UH and Biocenter Finland. In addition, we are grateful to Eija Korpelainen and Maria Lehtivaara in CSC for helping to run Chipster. Castrén lab was supported by the ERC grant # 322742 – iPLASTICITY, the Sigrid Jusélius foundation, Jane & Aatos Erkko Foundation, Academy of Finland grants #294710, # 327192 and #307416, EU Joint Programme - Neurodegenerative Disease Research (JPND) *CircProt* project co-funded by EU and Academy of Finland #643417, the doctoral program Brain&Mind and the bilateral exchange program between Academy of Finland and JSPS (Japan Society for the Promotion of Science).

## Author contributions

F.W., J.U. and E.C. conceived and designed the project. F.W. performed the experiments. M.V. analyzed the in vivo electrophysiology recordings under the supervision of S.P.. G.D. helped with the Western Blot experiments. M.L. cut the brains on the vibratome and helped with immunohistochemistry. E.J., S.M., and J.U. helped with single nuclei sequencing experiments and data analysis. A.S. helped with the behavioral experiment using hPV-TrkB CKO mice combined with fluoxetine treatment. J.H. analyzed the PV/PNN intensities. S.K., C.R., J.E., M.R., S.K., T.T. and S.L. provided supervision and support during the electrophysiological experiments. F.W., J.U. and E.C. wrote the manuscript.

## Material and Methods

### EXPERIMENTAL MODEL AND SUBJECT DETAILS

All animal procedures were done according to the guidelines of the National Institutes of Health Guide for the Care and Use of Laboratory Animals and were approved by the experimental Animal Ethical Committee of Southern Finland. Female mice (PV-cre, WT or hPV-TrkB CKO) older than 110 days were group housed with *ad libitum* access to food and water. For the shift in ocular dominance paradigm, the mice underwent transparent skull surgery and were monocularly deprived for 7 days.

#### Mice

Heterozygous mice with TrkB deletion specifically in PV^+^ interneurons (PV-hTrkB mice; PV^pvr/wt^, TrkB^flx/wt^) were produced by mating females of heterozygous PV specific Cre line (PV^pvr/wt^) (Pvalb-IRES-Cre, JAX: 008069, Jackson laboratory) (Hippenmeyer et al., 2005) and male of homozygous floxed TrkB mice (TrkB^flx/flx^) (Minichiello et al., 1999). Single floxed mice (TrkB^flx/wt^) served as control. For patch-clamp electrophysiology, females of the homozygous PV Cre line (PV^pvr/pvr^) were crossed with males harboring a homozygous TdTomato indicator allele (Rosa26^TdT/TdT^) (Ai14, JAX: 007914, Jackson laboratory, Bar Harbor, ME) (Madisen et al., 2010) and heterozygous floxed TrkB allele (TrkB^flx/wt^)(Minichiello et al., 1999) to reproduce compound transgenic mice with TrkB deletion and TdT expression specifically in PV interneurons (PV^pvr/wt^, TrkB^flx/wt^, Rosa26^TdT/wt^) (hPV-TrkB CKO) and wild-type allele in TrkB (PV^pvr/wt^, TrkB^wt/wt^, Rosa26^TdT/wt^) (WT). All of the parental strains were back-crossed with C57BL/6J for more than six generations. To assure the complete closure of the critical periods the mice were older than 110 days at the start of the experiments. The mice were kept under 12h light/dark cycle with light on at 6 am and *ad libitum* access to food and water.

#### Construction of optoTrkB

Photolyase homology region (PHR) domain of optoTrkB (Chang et al., 2014) was optimized in codon usage for mice and connected with a flexible tag (Kennedy et al., 2010) to the C-terminus of a full-length mouse TRKB, as previously described (Umemori et al, submitted). For cre-dependent expression of optoTrkB, a double floxed inverted open-reading frame (DIO) structure (Fenno et al., 2014) of optoTrkB (DIO-optoTrkB) was constructed (Figure S1A). We artificially synthesized DNA including human Synapsin promoter (Kügler et al., 2001), lox2272 (Lee & Saito, 1998), loxP, inverted optoTrkB sequence [PHR, flexible tag (Kennedy et al., 2010), mouse full-length TrkB (NM_001025074)], lox2272, loxP, and cloned into pFCK(0.4)GW (a gift from Pavel Osten) (Addgene plasmid # 27229; http://n2t.net/addgene:27229; RRID:Addgene_27229) (Dittgen et al., 2004) using PacI and EcoRI cloning sites (Figure S1a). Furthermore, for electrophysiological studies, we constructed DIO-optoTrkB expressing TdTomato (Shaner et al., 2004) (DIO-optoTrkB-IRES-TdT) (Figure S1C). A fragment of Internal ribosome entry site (IRES) with TdTomato was amplified by PCR with primers (5’- ggcgcgCCCCCCTCTCCCTCCCCCCC -3’ and 5’- ggcgcgccTTACTTGTACAGCTCGTCCATGCCGTACAG -3’) using Hot start Q5^®^ polymerase (NEB, Frankfurt, Germany) from LeGO-iT (a gift from Boris Fehse) (Addgene plasmid # 27361; http://n2t.net/addgene:27361; RRID:Addgene_27361) (Weber et al., 2010), and cloned into pCR™ Blunt II-TOPO^®^ vector (ThermoFisher Scientific, Hvidovre, Denmark). Then the sequence was confirmed by sequencing and sub-cloning into AscI cloing sites of DIO-optoTrkB (Figure S1A) to construct DIO-optoTrkB-IRES-TdT. Cre-dependent Inversions of DIO-optoTrkB and DIO-optoTrkB-IRES-TdT were confirmed in HEK293 cells co-transfected with the DIO-vectors and plasmids expressing Cre recombinase followed by PCR (Set1 (Figure S1B), and immunocytology (Figure S1D). The following primers were used to confirm the inversion of DIO-optoTrkB:

Set1 (5’- AGTCGTGTCGTGCCTGAGA -3’ and 5’- GAAATTTATGTGCCGCAGGT -3’); Set2 (5’- CTGCTGGCAAAGGCTATTTC -3’ and 5’- GGGCCACAACTCCTCATAAA -3’).

#### Virus generation

DIO-optoTrkB/DIO-optoTrkB-IRES-TdT, pLP1, pLP2, and pVSVG were co-transfected into HEL293FT cells by jetPEI^®^ (Polyplus Transfection, Illkirch, France) according to the manufacturer’s instruction, for producing lentivirus through a previously reported method (Hioki et al., 2007). Briefly, 3μg of the plasmids was transfected into HEL293FT cells and cultured on a poly-D-Lysine coated dish containing pre-warmed Opti-MEM^®^ medium (ThermoFisher Scientific) with 2% FBS for 72 hrs. Then the culture supernatant was collected and exchanged with a new Opti-MEM^®^ medium with 2% FBS followed by incubation for 72 hrs. The culture supernatant was collected again and centrifuged at 4 °C at 2000 g for 10 min. The supernatant was then concentrated with Amicon^®^ Centrifugal Filter Units Ultra −15 (Merck Millipore, Darmstadt, Germany) into less than 10ml. The concentrated solution was purified using the sucrose gradient method (Tiscornia et al., 2006) and aliquots were stored at −80 °C until use. From one aliquot the p24 capsid protein concentration was measured to estimate the infection unit (IU).

#### Virus injection

PV-cre mice were anaesthetized with 2.5% isoflurane and DIO-optoTrkB (p24 concentration/titer 4,04 × 10^7) was stereotaxically injected into the binocular area of the V1, previously identified through optical imaging (see below) and using blood vessels as landmarks. DIO-optoTrkB was stimulated with blue LED light (470nm) through the transparent skull during monocular deprivation twice a day for 30 seconds at 8-10 am and 4-6 pm. Blue LED light was provided by a BLS Super High Power Fiber-Coupled LED Light Sources (BLS-FCS-0470-100, Mightex, Pleasanton, CA) connected to a BioLED Light Source Control Modules (BLS-3000-2, Mightex). To avoid light exposure through the skull, the transparent skull was covered by black nail polish between light stimulation sessions.ht

#### Surgery

For chronic imaging of intrinsic signals, the animals underwent transparent skull surgery as described previously (Steinzeig et al., 2017).

#### Fluoxetine treatment

Fluoxetine (Bosche Scientific, New Brunswick, NJ) was administered via drinking water (0.08 mg/ml), corresponding to a dose of approximately 8-12 mg/kg/day, and kept in light-protected bottles. The drinking water of both the control and fluoxetine group contained 0.1 % saccharin, was changed twice a week and the consumption was monitored. Drug treatment started after transparent skull surgery and continued throughout the whole experiment.

#### Monocular deprivation and virus stimulation

The animals were anaesthetized with intraperitoneal injection as described above. The eye lashes were trimmed and the eye lid margins were sutured shut with 3 mattress sutures. To prevent postoperative infections, an eye ointment containing dexamethasone was applied. The eyes were checked daily until reopening, re-sutured if needed, and mice with signs of corneal injury were excluded from the experiments.

#### Optical imaging of intrinsic signals

We determined the strength of neuronal responses to stimulation of either eye in the binocular region of the V1 using imaging of intrinsic signals (IOS) (Kalatsky & Stryker, 2003) before (IOS I; IOS III) and after (IOS II; IOS IV) monocular deprivation. The animals were chamber anaesthetized with 1.8% isoflurane in a 1:2 mixture of O_2_/air and then intubated and ventilated with 1.2% isoflurane in the same mixture.

Intrinsic optical signal responses were recorded from the V1 of the right hemisphere according to a previously described protocol (Kalatsky & Stryker, 2003), which was modified for the measurement of OD plasticity (Cang et al., 2005).

#### Data analysis of optical imaging

Cortical maps were computed based on the acquired frames using Fourier decomposition to extract the signal from biological noise using an analysis software package for continuous recording of optical intrinsic signals (VK Imaging, USA) (Kalatsky & Stryker, 2003). The ODI was then calculated for every pixel within the binocularly responding region based on the formula (C−I)/(C+I), where “C” refers to the response magnitude of the contralateral eye and “I” to that of the ipsilateral eye. For each animal, several ODIs were collected and then averaged. Positive ODI values represent contralateral dominance, negative represent ipsilateral dominance, while ODI values of 0 correspond to equally strong contralateral and ipsilateral eyes.

#### In vivo electrophysiology

To measure oscillations in the binocular region of the optoTrkB transfected area, PV-cre mice underwent transparent skull surgery and optoTrkB was injected as described above. Under 15% urethane anesthesia (in PBS), a hole for a 16 channel optrode (A1×16-10mm-100-177-OA16LP, NeuroNexus, Ann Arbor, MI) was drilled next to the injection site and a hole for a ground electrode next to lambda. The animals were head-fixed and the optrode was covered with DiI and slowly inserted into the V1 to a depth of about 2200 μm. The data was recorded using a Smartbox (NeuroNexus) at a sampling rate of 20 kHz. After collecting a 20 minute baseline, optoTrkB was stimulated for 30 seconds with blue LED light and recorded for 2h, separated into 20-minute blocks. Finally, the animals were transcardially perfused and the brains fixed with 4% PFA. To confirm the position of the electrodes and co-localization with optoTrkB, 250 μm coronal slices were cut on a vibratome.

The digitized data were first offline notch filtered at 50 Hz by the Python RHD2000 interface provided by Intan. Afterwards, the data were band-pass filtered between 4 and 150 Hz, using a 2nd order Butterworth filter with zero-phase shift, and then downsampled to a sampling rate of 1 kHz to extract the local field potential (LFP) component. The LFP for each electrode contact was normalized to zero and the underlying current source density (CSD) was calculated as the second spatial derivative, using previously published methods (Voigt & Kral, 2019).

Time-frequency analysis of the CSD data was performed by wavelet filtering using 36 Morlet wavelets with a width parameter m = 6 and with frequencies ranging from 4 to 112 Hz with log-constant scaling. For further analyses, the signal from all electrodes that were identified to reside in areas with optoTrkB expression by the histological analysis was averaged, and the data were binned into non-overlapping 20 seconds bins. Morlet power spectra were estimated by computing the averaged magnitude over all bins of the 20-minute recording period per condition. For statistical analysis, broadband power in the 20-minute baseline period and the 20-minute period beginning 100 minutes after LED onset was compared using permutation testing. Surrogate data was generated by 1000 random permutations of condition labels and the effect size was measured using Cohen’s d. All data analyses and statistics were performed in Python (ver. 2.7) using SciPy (ver. 0.19.0), the NeuroDSP toolbox (ver. 2.0.1-dev)(Cole et al., 2019), and custom written scripts.

#### Electrophysiology in acute slices

The brains of optoTrkB-infected PV-cre mice were dissected in darkness using red light illumination, and kept in the dark throughout the whole experiment. The brains, including brains of fluoxetine-treated WT and hPV-TrkB cKO mice, were dissected and immersed in ice-cold dissection solution containing (in mM): 124 NaCl, 3 KCl, 1.25 NaH_2_PO4, 1 MgSO_4_, 26 NaHCO_3_, 15 D-glucose, 9 MgSO_4_ and 0.5 CaCl_2_. The cerebellum and anterior part of the brain were removed and coronal 350μm brain slices of the V1 were cut on a vibratome (Leica Biosystems, Wetzlar, Germany). Slices were divided into two groups and allowed to recover for 30 min at 31-32°C in artificial cerebrospinal fluid (ACSF) containing (in mM): 124 NaCl, 3 KCl, 1.25 NaH_2_PO_4_, 1 MgSO_4_, 26 NaHCO_3_, 15 D-glucose, and 2 CaCl_2_ and bubbled with 5% CO_2_/95% O_2_.

OptoTrkB-transfected slices in one of the groups were acutely stimulated for 30 seconds with blue light after transferring them to the recording chamber, whereas the transfected slices in the other group were kept in darkness.

Field excitatory postsynaptic currents (fEPSPs) were recorded in an interface chamber (32°C) with ACSF-filled glass microelectrodes (2-4 MΩ) positioned within layer II/III of the V1 using an Axopatch 200B amplifier (Molecular devices, San Jose, CA). Electric stimulation (100 μsec duration) was delivered with a bipolar stimulation electrode placed at the border of the white matter (WM) and layer VI. Baseline synaptic responses were evoked every 20 seconds with a stimulation intensity that yielded a half-maximum response. After obtaining a 15 minute stable baseline, θ burst stimulation (TBS) (4 sweeps at 0.1 Hz, each sweep with 10 trains of 4 pulses at 100 Hz at 200 ms intervals) was delivered and field potentials in response to 0.05 Hz stimulation were recorded for additional 45 minute. WinLTP (0.95b or 0.96, www.winltp.com) was used for data acquisition and analysis.

To measure intrinsic excitability, the brains of PV-cre mice transfected with the DIO-optoTrkB-IRES-TdT lentivirus were dissected as described above but cut in a protective NMDG ACSF (Ting et al., 2014) containing (in mM): 92 NMDG, 2.5 KCl, 1.25 NaH_2_PO_4_, 30 NaHCO_3_, 20 HEPES, 25 glucose, 2 thiourea, 5 Na-ascorbate, 3 Na-pyruvate, 0.5 CaCl_2_·4H_2_O and 10 MgSO_4_·7H_2_O, pH 7.3–7.4 The slices were allowed to recover at 32°C for 10 min, after which they were transferred to modified ACSF containing (mM): 92 NaCl, 2.5 KCl, 1.25 NaH_2_PO_4_, 30 NaHCO_3_, 20 HEPES, 25 glucose, 2 thiourea, 5 Na-ascorbate, 3 Na-pyruvate, 2 CaCl_2_·4H_2_O and 2 MgSO_4_·7H_2_O, pH 7.3-7.4 for storage. Recordings were done in submerged chamber perfused with normal ACSF (32°C).

Whole cell patch clamp recordings from the TdT expressing PV cells were obtained under visual guidance under ambient light with glass microelectrodes (3-5 MΩ) filled with a low Cl^−^-filling solution containing (in mM): 135 K-gluconate, 10 HEPES, 2 KCl, 2, Ca(OH)_2_, 5 EGTA, 4 Mg-ATP, 0.5 Na-GTP using a Multiclamp 700A amplifier (Axon Instruments, Foster City, CA). Uncompensated series resistance (Rs) was monitored by measuring the peak amplitude of the current response to a 5 mV step. Only experiments where Rs < 30 MΩ, and with < 20 % change in Rs during the experiment, were included in analysis.

Intrinsic excitability was measured in current clamp mode by injecting currents ranging from −50 to 600 pA for 600 ms in 50 pA steps from the resting membrane potential of −60/−70 mV. The recordings were analyzed in Clampfit (Molecular Devices, San Jose, CA) programs. A minimum of three action potentials (AP) were averaged based on their AP half-width (10^th^ AP at 200pA injected from rheobase). sEPSCs were recorded under voltage clamp at −70 mV and analyzed in Clampfit (Molecular Devices). The threshold for detection of inward sEPSC events was three times the baseline noise level, and all detected events were verified visually.

#### Sample collection

For collecting tissue samples for qPCR and Western Blot experiments, the brains of optoTrkB infected PV-cre mice were dissected and immersed in ice-cold dissection solution containing (in mM): 124 NaCl, 3 KCl, 1.25 NaH_2_PO_4_, 1 MgSO_4_, 26 NaHCO_3_, 15 D-glucose, 9 MgSO_4_ and 0.5 CaCl_2_. The V1 was dissected and incubated at 31-32°C in artificial cerebrospinal fluid (ACSF) containing (in mM): 124 NaCl, 3 KCl, 1.25 NaH_2_PO_4_, 1 MgSO_4_, 26 NaHCO_3_, 15 D-glucose, and 2 CaCl_2_ and bubbled with 5% CO_2_/95% O_2_. The tissue samples were either immediately collected in NP lysis buffer and homogenized (control), or stimulated with blue LED light for 30 seconds and collected and homogenized after 15 minutes, 30 minutes or 60 minutes. All of the procedures were done in dark conditions. The samples were further used for qPCR and Western Blot analysis.

#### Western Blot

The samples were centrifuged (16000 g, 15 min at +4°C) and the supernatant was used to measure the protein concentrations using the Lowry method (Biorad DC protein assay, Bio-Rad, Richmond, CA) (Lowry et al, 1951). The samples were separated in a SDS-PAGE (2-4% gradient gel, NuPage™; ThermoFisher Scientific) and blotted to a PVDF membrane (300 mA, 1 h, + 41°C). The membranes were washed in Tris Buffer Solution with 0,001% Tween ^®^20 (TBST), blocked in TBST with 3% BSA for 1 hour at room temperature and incubated in primary antibody solutions (in TBST with 3% BSA) directed against: phosphorylated (pY816, #4168, 1:1000; pY515, #9141, 1:1000; pY706-7, #4621S, 1:1000, Cell signaling technology (CST), Danvers, MA) and non-phosphorylated forms of TrkB and CREB (TrkB, 1:2000, BD Transduction Laboratories, San Jose, CA; CREB, #4820S, CST, Danvers, MA) at +4°C for overnight. After washing in TBST, the membranes were further incubated in secondary antibody solutions (TBST with 5% Non-Fat Dry Skinned Milk and Horseradish Peroxidase conjugated secondary antibodies Goat Anti-Rabbit/Mouse, 1:10000, Bio-Rad) for 2 hours at room temperature. After washing with TBST and rinsing with PBS, secondary antibodies were visualized by an electrochemiluminescence kit (ECL plus, ThermoFisher Scientific) according to the manufactures instruction, and detected using a FUJIFILM LAS-3000 dark box.

#### qPCR

RNA was purified from the lysate following the manufacturer’s protocol using a combined protocol of QIAzol^®^ (Qiagen, Hilden, Germany) and NucleoSpin^®^ (Macherey-Nagel, Düren, Germany). Briefly, the aqua layer was isolated after Qiazol and chloroform extraction. The RNA was washed in 100% ethanol and the DNA was digested in the spin columns. The purified RNA was then reverse transcribed to cDNA using Maxima First Strand cDNA Synthesis Kit (ThermoFisher Scientific). The amount of cDNA synthesized from the target mRNA was quantified by real-time PCR (qPCR). The following primers were used to amplify specific cDNA regions of the transcripts of interest: Kv3.1 (5’ -AGAGATTGGCACTCAGTGACT-3’ and 5’ -TTGTTCACGATGGGGTTGAAG-3’), Kv3.2 (5’ -AGGCTATGGGGATATGTACCC-3’ and 5’ -TGCAAAATGTAGGCGAGCTTG-3’),PV (5’- TGTCGATGACAGACGTGCTC-3’ and 5’-TTCTTCAACCCCAATCTTGC-3’).

#### Immunohistochemistry

Animals were transcardially perfused with PBS followed by chilled 4% paraformaldehyde (PFA) in PBS. Brains were removed under ambient light and left for fixation in 4% PFA overnight at +4 °C. For cutting, the brains were embedded in 3% agar and cut into 40 μm coronal visual cortical sections using a vibratome (Leica Biosystems, Wetzlar, Germany). After washing with PBST (1x PBS and 0.2% TritonX100), the sections were incubated in 10% donkey serum (Vector Laboratories, Peterborough, UK) and 3% Bovine Serum Albumin (BSA) (Sigma-Aldrich, Steinheim, Germany) in PBST for 30 minutes at room temperature. Next, the sections were incubated with the following antibodies: 1) guinea pig anti-parvalbumin (1:1000) (#195004, Synaptic Systems, Göttingen, Germany), 2) biotinylated lectin from Wisteria floribunda (WFA; 1:200) (#L1516-2MG, Sigma-Aldrich), and 3) rabbit anti-phospho Kv3.1 (1:100) (#75-041, Phosphosolutions, Aurora, CO) overnight at +4° C. After washing in PBST, the samples were further incubated in: 1) goat anti-guinea pig secondary antibody conjugated with Alexa Fluor647/546 (1:1000) (Abcam, Cambridge, UK), 2) streptavidin conjugated with Alexa Fluor488 (1:1000) (Thermofisher Science), or 3) Goat anti-rabbit conjugated with Alexa Fluor 647 (1:1000) (Life technologies, Carlsbad, CA) for 2 hours at room temperature protected from light. After final washing in PBS, the sections were transferred to 0.1M PB with gelatin, mounted on glass slides and covered with DAKO mounting medium (Sigma Aldrich).

#### Image acquisition and analysis

Quantitative analysis of immunostainings was performed blind. Images were taken from the V1 according to the mouse brain atlas.

Laser scanning confocal microscopy was used to detect PV positive (PV^+^), PNN positive (PNN^+^), double positive (PV^+^PNN^+^) and pKv3.1-positive cells. Images were obtained using a confocal microscope LSM 700 (Carl Zeiss, Vantaa, Finland) equipped with a 10× objective lens (10x Plan-Apochromat 10x/0.45, Carl Zeiss) and imaging Software ZEN 2012 lite (Carl Zeiss). From each section, a z-stack containing at least 10 consecutive images was obtained. A minimum number of 3 sections per animal were imaged using the same microscope and the same camera settings for all samples. Image processing was done using ImageJ software version 1 (Schneider et al., 2012). All images in each z-stack were analyzed and the number of cells was averaged per z-stack.

To determine the PV cell populations, frequency distribution analyses were performed on PV intensities taken from non-stimulated optoTrkB samples or control WT samples to serve as reference group. The PV cell populations were defined as low PV (0-8000 a.u.), intermediate-low PV (int-low PV, 8000-16000 a.u.), intermediate-high PV (int-high, 16000-24000 a.u.) and high PV (24000-36000 a.u.) expressing cells and these criteria were applied to the light stimulated samples.

#### Nuclei preparation

Single nuclei RNA sequencing (snRNA-seq) was conducted by following previously reported methods with some modifications (Bakken et al., 2018; Krishnaswami et al., 2016) (Figure S3A). Considering that optoTrkB is sensitive to light stimulation, we used snRNA-seq instead of single cell RNA sequencing (scRNA-seq) after FACS selected by TdTomato fluorescence, to avoid laser-induced optoTrkB activation during FACS. There is a good correlation of gene expression and gene detection sensitivity in each cell between SnRNA-seq and scRNA-seq (Bakken et al., 2018), and snRNA-seq has several advantages, such as reduced dissociation bias and dissociation-induced transcriptional stress responses (Wu et al., 2019). Tissue samples were obtained bilaterally from the visual cortex of adult female PV-cre mice infected with DIO-optoTrkB-TdT. After deep anesthesia with pentobarbital, the mice were perfused with ice-cold NMDG ACSF (Ting et al., 2014) consisted of 0.5 mM CaCl_2_, 25mM D-Glucose, 98 mM HCl, 20 mM HEPES, 10 mM MgSO_4_, 1.25 mM NaH2PO4, 3 mM Myo-inositol, 12 mM N-acetylcysteine, 96 mM N-methyl-D-glucamine, 2.5 mM KCl, 25 mM NaHCO_3_, 5 mM sodium L-Ascorbate, 3 mM sodium pyruvate, 0.01 mM Taurine, and 2 mM Thiourea, bubbled with a carbogen gas (95% O_2_ and 5% CO_2)_, and the brains were isolated and sliced on a vibratome (MICROM, HM 650V, Thermofisher Science) in NMDG ACSF (Ting et al., 2014). Then, the slices were exposed to blue LED light (30 seconds), and incubated in a modified HEPES ACSF including 92 mM NaCl, 2.5 mM KCl, 1.25 mM NaH_2_PO_4_, 30 mM NaHCO_3_, 20 mM HEPES, 25 mM glucose, 2 mM thiourea, 5 mM Na-ascorbate, 3 mM Na-pyruvate, 2 mM CaCl_2_, 2 mM MgSO_4,_ 3 mM Myo-inositol, and 0.01 mM Taurine for one hour. The visual cortex was bilaterally isolated from the slices under a microscope using red light illumination, and tissues were transferred to pre-cooled Dounce homogenizers filled with cold homogenization buffer (250 mM sucrose, 25 mM KCl, 5 mM MgCl2, 10 mM Tris buffer, pH 8.0, 1 μM DTT, 1× protease inhibitor (cOmplete, Sigma-aldrich), 0.4 U/μl RNase Inhibitor (Promega), 0.2 U/μl Superasin (Thermofisher Science, Waltham, MA), 0.1% (v/v) Triton X-100 (Sigma-Aldrich), 10 μg/ml Cyclohexamide (Sigma-Aldrich), 10 μg/ml Actinomycin D (Sigma-Aldrich), 10 ng/ml Hoechst 33342 (Thermofisher Science)). The tissues were homogenized with five strokes of the loose pestle, followed by 10 strokes of the tight pestle. Finally, nuclei of the tissues were obtained by filtration through a BD Falcon™ cell strainers (BD Falcon, San Jose, CA). All procedures were performed with RNase free in the dark condition.

#### Immunostaining and sorting of nuclei

The nuclei were centrifuged (1,000 x g, 8 min, 4 °C) and resuspended in staining buffer (PBS, pH 7.4, with 0.5% RNase-free BSA (Promega, Heidelberg, Germany) and 0.2 U/μl of RNasin Plus RNase inhibitor (Promega)). Samples were incubated in blocking/washing buffer (PBS with 0.5% BSA). The nuclei were incubated with mouse monoclonal anti-NeuN antibody (1:5000) (MAB377, Sigma-Aldrich) or purified mouse IgG1k (1:5000) (554121, BD Pharmigen, San Diego, CA) as a negative control for staining and sorting for 30 min with rotation at 4 °C. Then the samples were washed in staining buffer by inverting the tubes several times, and were centrifuged for 5 min at 400g at 4 °C. The pellet nuclei were re-suspended in 500 μl of staining buffer containing goat anti-mouse Alexa Fluor 488 secondary antibody (Thermofisher Science) with a final dilution of 1:5000, and were incubated with rotation for 30 min at 4 °C followed by washing with blocking/washing buffer.

#### FACS of nuclei

Hoechst 33342 and NeuN positive nuclei stained with Alexa488 were sorted and collected by BD Influx cell sorter (BD Biosciences, Heidelberg, Germany) (Figure S3B). After checking the quality, collected nuclei were loaded on Chromium Controller (10x Genomics).

#### Single nuclei RNA sequencing (snRNA-seq)

Preparation of cDNA Library was done using Chromium Single Cell 3’ Gene Expression v3 library preparation kit (10x Genomics), and sequenced with NovaSeq 6000 (Illumina, San Diego, CA) with read lengths: 28bp (Read 1), 8bp (i7 Index), 0 bp (i5 Index) and 89bp (Read 2). Data was pre-processed using CellRanger 3.0. The reads are mapped to reference genome that included introns, and we obtained 3,789 and 6,923 cells with approximately 10k median reads per cell (Figure S4).

#### Data analysis for snRNA-seq

Data analysis was carried out using Chipster version 3.16 (Kallio et al., 2011), which utilizes Seurat v3 (Butler et al., 2018). Cells with more than 7,000 and less than 100 reads were filtered out, and reads for the remaining cells were scaled logarithmically. Based on results for plotting variance against expression, 3000 most highly varied genes in 10,706 cells were used for clustering and sample integration. By Seurat’s integrated analysis (Butler et al., 2018), 20 canonical components were used for finding anchors and 20 principal components to integrate the samples together. Clusters were obtained using UMAP (Mcinnes et al., 2018) with 20 principal components and resolution 0.5 (Figure S5A, and S5B). This resulted in 18 clusters and we identified the clusters of pyramidal neurons and interneurons with previously reported markers (Tasic et al., 2018), such as *Glutamate decarboxylase 1* (*Gad1),* and those of interneurons with *Solute Carrier Family 6 Member 1* (*Slc6a1)*, *Parvalbumi*n (*Pvalb*), *Paired box protein* (*Pax6*), *Vasoactive intestinal peptide* (*VIP*), and *Somatostatin* (*Sst*) (Fig. 2A) (Figure S5C). Then a subset of data containing only parvalbumin-positive interneurons was selected and clustered again by UMAP with the same parameters resulting in four different clusters (Fig. 2B) (Figure S6A and S6B). Markers and differentially expressed genes between non- and light-exposed samples were detected in each cluster (Figure S6D) (Table S3–6). Pathway and gene ontology analysis was carried out with DAVID (Huang et al., 2009).

#### Statistical analysis

All statistical graphs were generated using Graphpad Prism v.6.07 (GraphPad Software, San Diego, CA). Unpaired t-Test, Two-way and one-way ANOVA followed by Tukey’s or Bonferroni post hoc tests were also performed using Graphpad Prism v.6.07. The confidence level was set to 0.05 (P value) and all results were presented as means±s.e.m. All the individual data points are shown in the histograms. The data distribution was assumed to be normal but this was not formally tested. The sample size was determined based on our previous experience.

### DATA AND SOFTWARE AVAILABILITY

snRNA-seq data have been deposited in NCBI’s Gene Expression Omnibus (Edgar et al., 2002) and are accessible through GEO Series accession number GSE 142797 (https://www.ncbi.nlm.nih.gov/geo/query/acc.cgi?acc=GSE142797). Other raw data is available in Mendeley Data, V1, doi: 10.17632/992cg2vrj5.1 (https://data.mendeley.com/datasets/992cg2vrj5/draft?a=ab1dd775-63b5-4eca-aa01-0a565fb6fa0d).

## Supplemental Information

**Figure S1.**
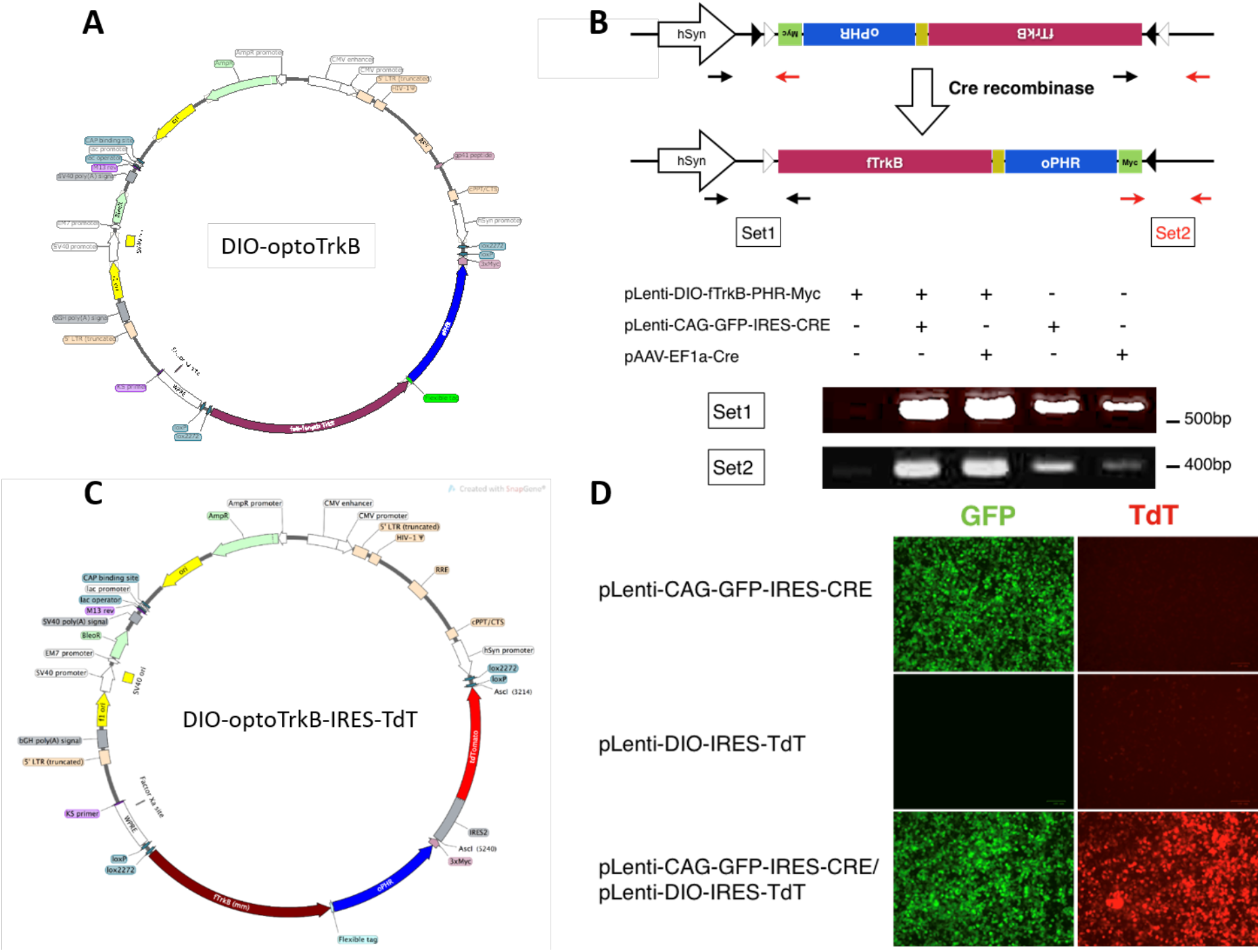
Confirmation of inversion of DIO-optoTrkB by cre-recombinase. (A) The plasmid producing lentivirus DIO-optoTrkB. (B) Confirmation of inversion of DIO-optoTrkB by cre-recombinase. HEK293 cells were co-transfected with pLenti-DIO-optoTrkB and pLenti-CAG-GFP-IRES-CRE or pAAV-EF1a-Cre, which express cre-recombinase. PCR analysis revealed that DIO-optoTrkB was inverted after co-transfection with cre but not when expressed alone. (C) The plasmid producing DIO-optoTrkB-IRES-TdTomato. (D) Confirmation of inversion of DIO-optoTrkB-IRES-TdTomato by cre-recombinase. HEK293 cells were transfected with either or both pLenti-CAG-GFP-IRES-CRE and pLenti-optoTrkB-IRES-TdTomato. Only co-transfected cells expressed TdTomato demonstrating that the expression is cre-dependent. hSyn, human Synapsin promoter; oPHR, optimised Photolyase homology region; fTrkB, full-length TrkB.

**Figure S2.**
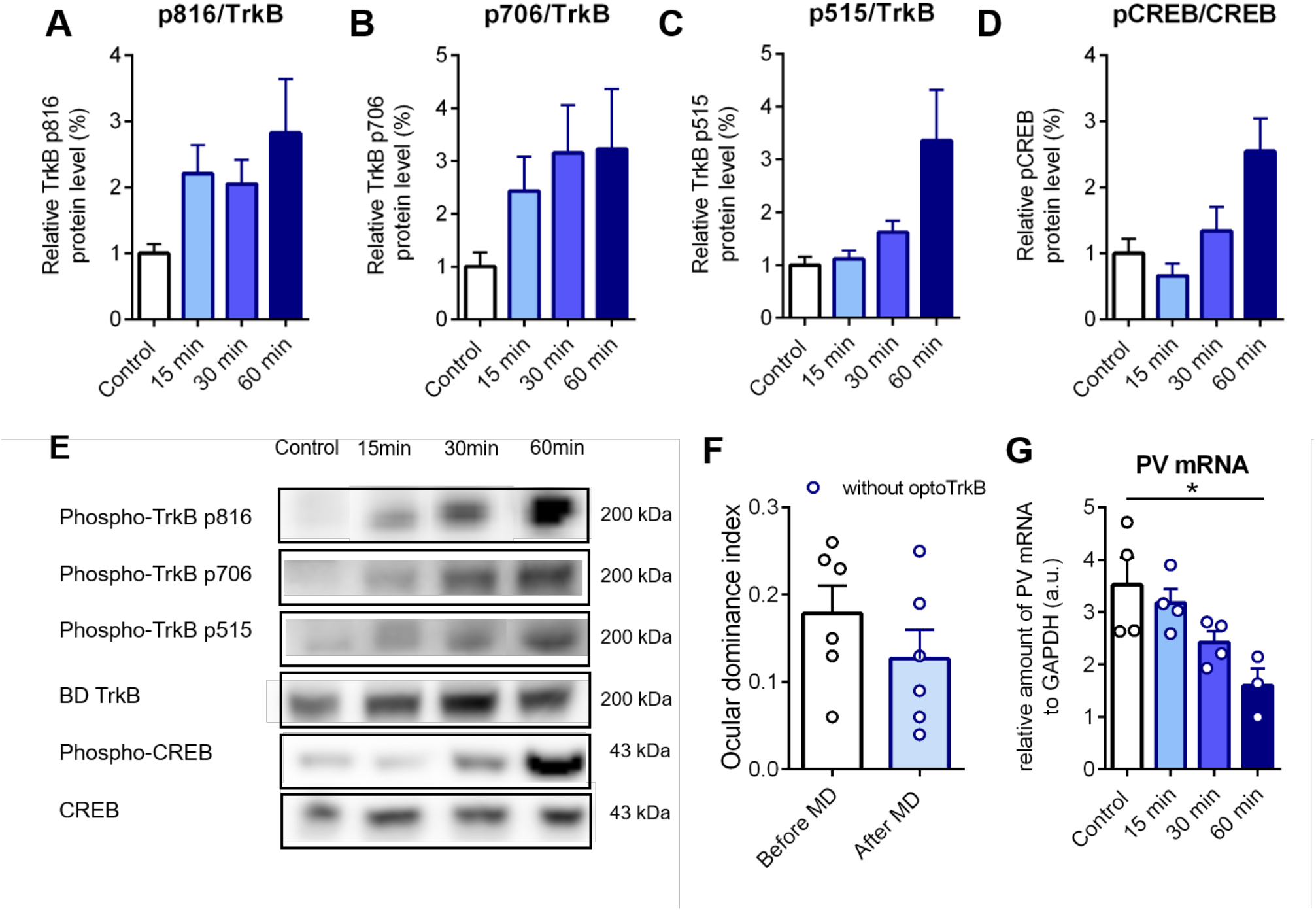
OptoTroB activation mediates plasticity in the visual cortex. (A-E) Western Blot analysis of V1 of PV-cre mice infected with DIO-optoTrkB without light stimulation (Control) and 15 minutes, 30 minutes and 60 minutes after light stimulation. Acute stimulation of optoTrkB with light results in increased phosphorylation of tyrosine sites: (A) Y816, (B) Y515, (C) Y706 and (D) phosphorylation of CREB. (E) Representative Western Blots of Y816, Y515, Y706 and CREB. (F) PV-cre mice without optoTrkB expression stimulated with blue light twice daily during 7 days of MD show no shift in ocular dominance. (G) PV mRNA levels are decreased 60 min after light stimulation measured by qPCR. qPCR measurements of control, 15 minutes, 30 minutes and 60 minutes after light stimulation. OptoTrkB reduces the expression of PV 60 minutes after stimulation. One-way ANOVA (F (3, 11) = 5.112; p = 0.0186) with Bonferroni’s post-hoc test comparing control vs. 15 min, 30 min and 60 min (control vs. 15 min, p = > 0.9999; control vs. 30 min, p = 0.1456; control vs. 60 min, p = 0.0125). n = 3-4 animals/group. Bars represent means ± SEM. * p < 0.05

**Figure S3.**
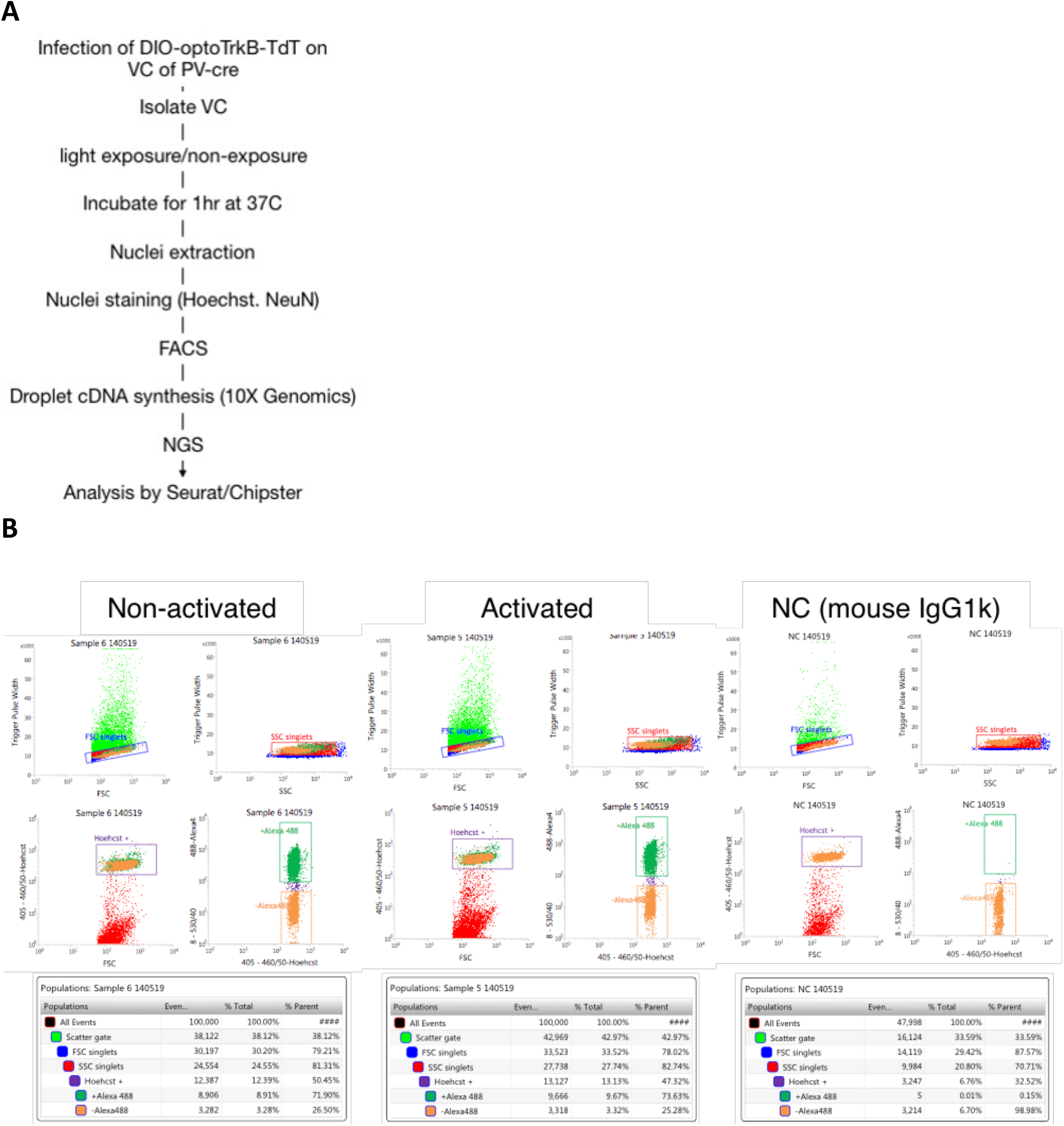
snRNA-seq after activation of optoTrkB. (A) Scheme of snRNA-seq. The primary visual cortices were bilaterally isolated from PV mice transfected by DIO-optoTrkB-TdT. Nuclei were extracted from the tissue, and Hoechst+ and NeuN+ cells were sorted by FACS. A cDNA library was constructed with Poly (A)-RNA in droplets. The fragmented cDNA, were then sequenced on a Next generation sequencer (NGS). (B) Sorting nuclei of neurons by FACS. Cells were stained with Hoechst 33342, NeuN antibody, and mouse IgG antibody (negative control) followed by labelling with anti-mouse IgG conjugated with Alexa488, and then sorted by BD Influx cell sorter (BD Biosciences). Hoechst 33342 positive nuclei were efficiently sorted in all samples. While small portion of nuclei (0.15%) labelled with IgG bound to Alexa488 were sorted in negative control, large portion of nuclei (∼72%) stained with NeuN bound to Alexa488 were efficiently sorted and collected.

**Figure S4.**
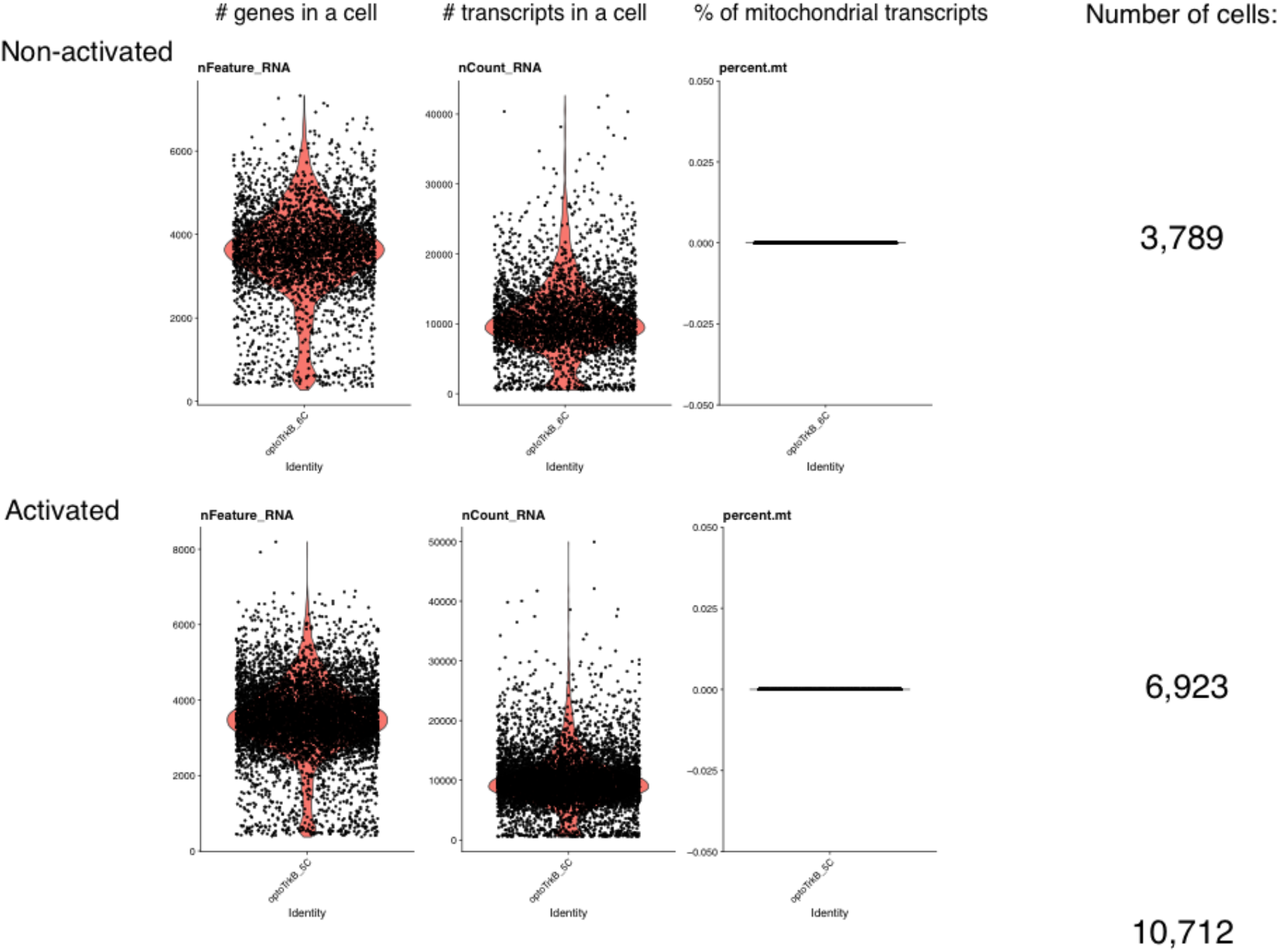
Number of genes, reads, and mitochondrial genes in nuclei. The plots represent genes and transcripts per cell, where the horizontal axis is cell id and vertical axis is the number of genes or transcripts, respectively. Each dot is a unique cell and orange areas represent cell density at vertical axis. In the right panels, the horizontal axis is cell id and the vertical axis is the percentage of transcripts originating from mitochondrial DNA. Each cell should be represented with a dot and the black line in 0.000 % shows absence of mitochondrial genes. Since we used only nuclei, mitochondrial genes are absent. Based on genes and transcripts per cell, we decided to remove cells that had fewer than 100 or greater than 7000 reads.

**Figure S5.**
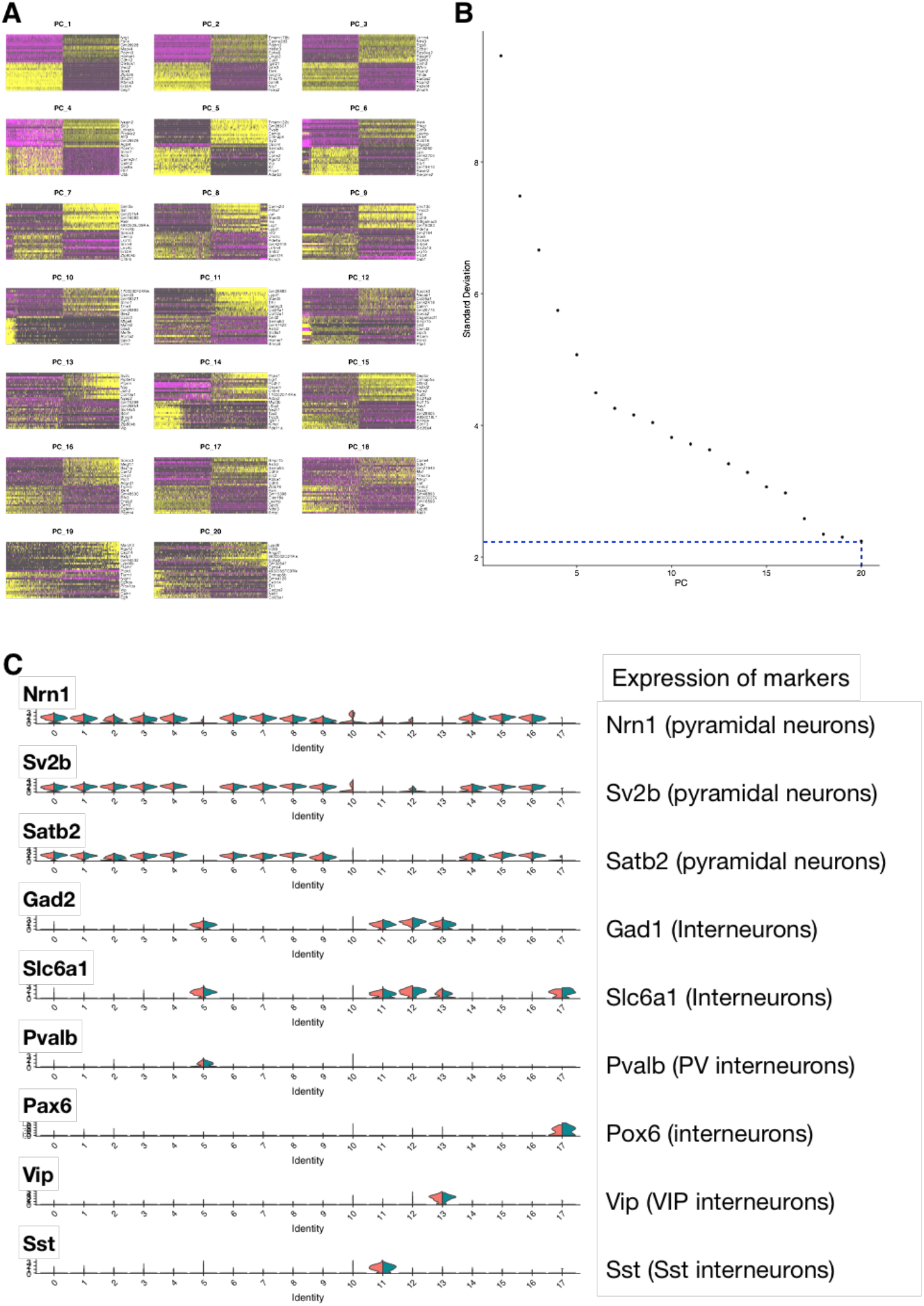
Clustering single nuclei. (A) For 20 first principal components, heatmaps are drawn with cells as columns and the 15 most important genes as rows. The heatmaps show major variation captured in 20 first principal components. (B) Elbow plot is generated by calculating how much of the variation is explained by the first 20 principal components. Horizontal axis represents ordinal number of principal component and vertical axis the standard deviation of explained percentage in variation. Most of the variation in the data is included in 20 first principal components, which were used for later analysis. (C)In the violin plot, the vertical axis indicates the expression level of a gene in a cluster (number below), and the horizontal axis indicates cell density at a given expression level. Pyramidal neurons were identified by using the markers Nrn1, Sv2b and Satb2 and while interneurons were done by Gad1 and Slc6a1. The cluster 5, 11, and 13 contained parvalbumin-, somatostatin-, and VIP-positive interneurons, respectively.

**Figure S6.**
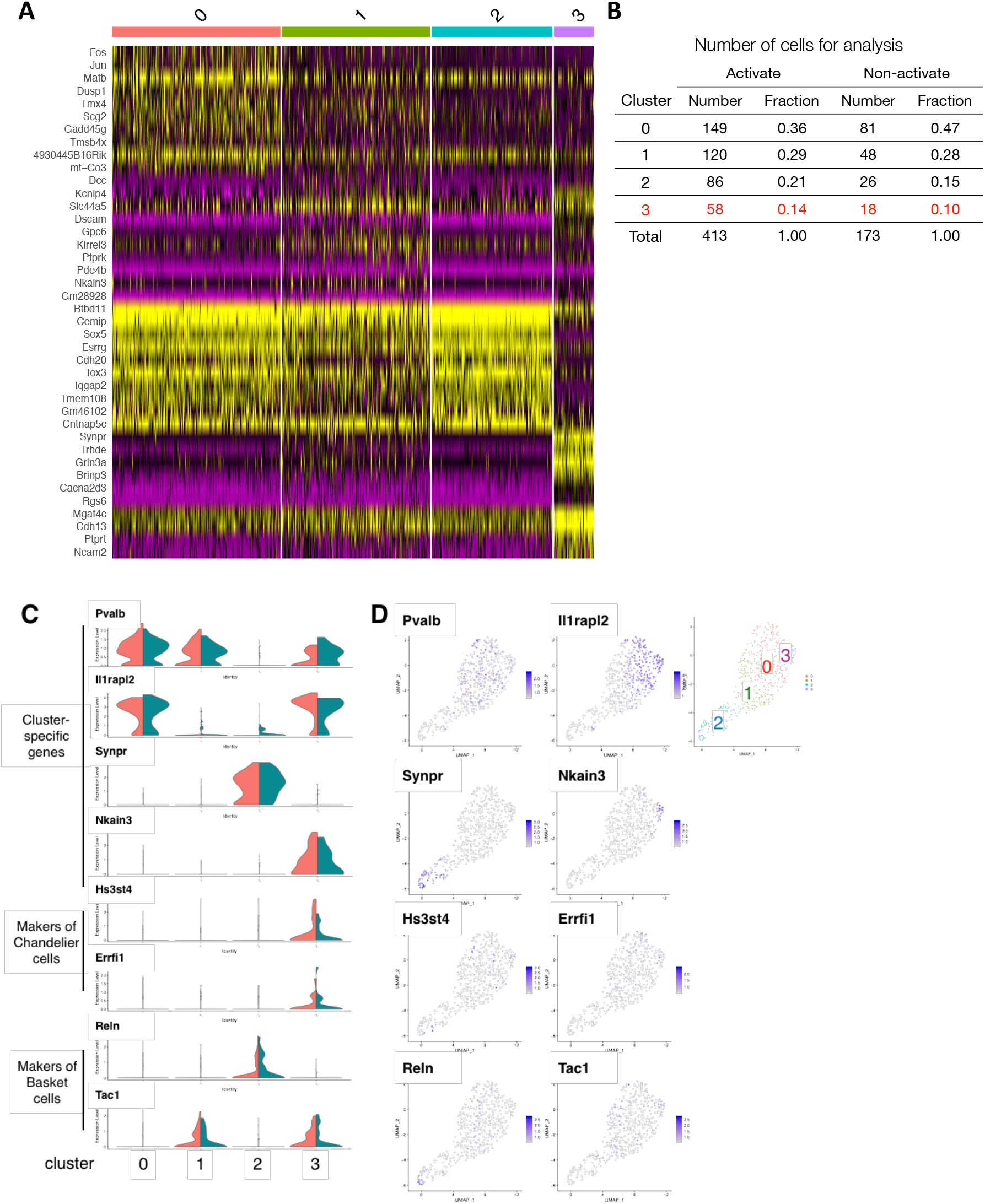
Clustering PV interneuron. (A) Genes in representative PCAs explain four clusters in PV neurons. Heatmaps of unique genes in each clusters show that the PV cluster can be further divided into four clusters (0-3). (B) Number and ratio of PV cells used for clustering and DE analysis. (C) Distribution of expression of genes in cluster specific genes (*Pvalb, Il1rapl2, Synpr, Nkain3*), representative markers for chandelier cells (*Hs3st4, Errfi1*), and basket cells (*Reln, Tac1*) as reported previously(Tasic et al., 2018). (D) Distribution of cells expressing each marker in the four clusters of PV interneurons. These results demonstrate that each cluster has unique expression of marker genes, and the cluster 3 includes both chandelier and basket cells.

**Figure S7.**
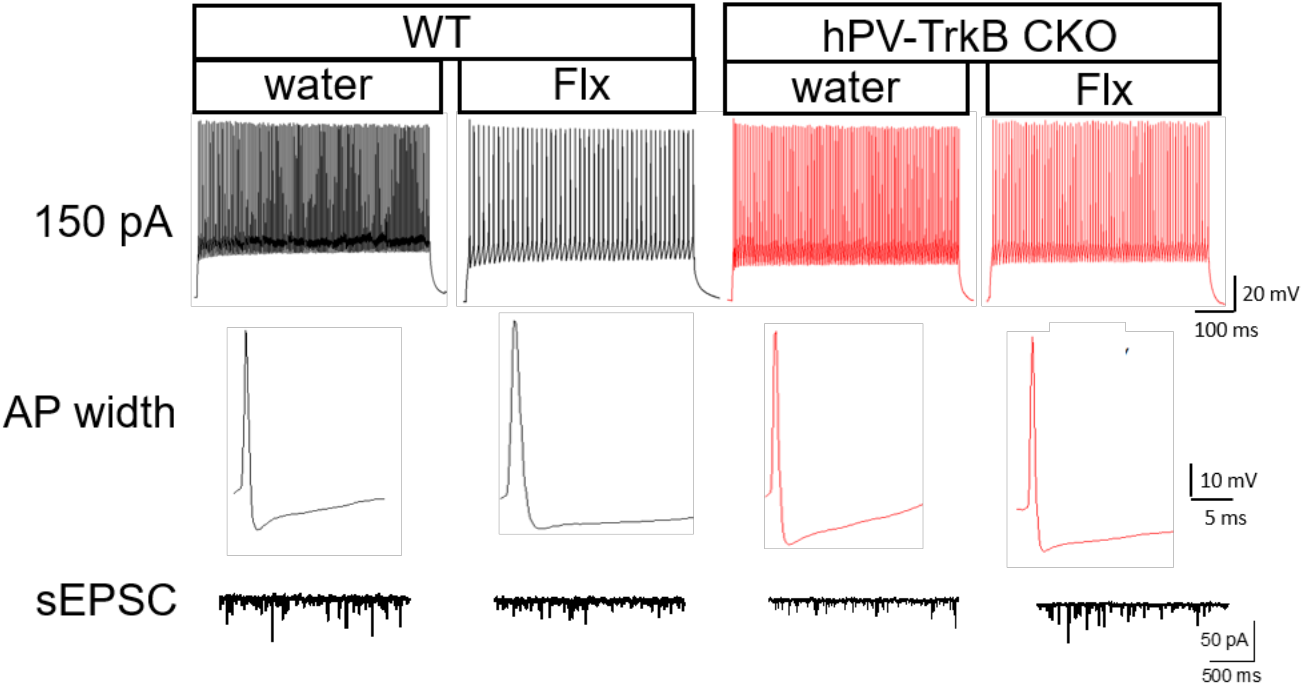
Representative traces of electrophysiological recordings. Representative traces used to estimate intrinsic excitability (150 pA, top row), AP half-width (2nd row) and sEPSC (bottom row) in WT mice treated with water (first column), WT mice treated with Flx (2nd column), CKO mice treated with water (3rd column) and CKO mice treated with Flx (last column).

**Table S1.** Number of nuclei in each cluster. (a) All clusters were identified by expression of makers previously reported (Tasic et al., 2018). (b) Number of cells in each cluster were provided by Chipster/Seurat version 3. (c) Percentage was calculated by dividing the number of nuclei in each cluster with that of total nuclei in non-activated or activated sample

| Cluster | Neuronal type<sup>a</sup> | Non-activated |  | Activated |  |
| --- | --- | --- | --- | --- | --- |
|  |  | N<sup>b</sup> | %<sup>c</sup> | N<sup>b</sup> | %<sup>c</sup> |
| 0 | Pyramidal | 696 | 18.4 | 1200 | 17.3 |
| 1 | Pyramidal | 349 | 9.2 | 874 | 12.6 |
| 2 | Pyramidal | 368 | 9.7 | 827 | 11.9 |
| 3 | Pyramidal | 390 | 10.3 | 746 | 10.8 |
| 4 | Pyramidal | 222 | 5.9 | 408 | 5.9 |
| 5 | Internuron | 173 | 4.6 | 413 | 6.0 |
| 6 | Pyramidal | 218 | 5.8 | 360 | 5.2 |
| 7 | Pyramidal | 172 | 4.5 | 334 | 4.8 |
| 8 | Pyramidal | 261 | 6.9 | 235 | 3.4 |
| 9 | Pyramidal | 192 | 5.1 | 293 | 4.2 |
| 10 | Pyramidal | 189 | 5.0 | 266 | 3.8 |
| 11 | Internuron | 146 | 3.9 | 237 | 3.4 |
| 12 | Internuron | 82 | 2.2 | 169 | 2.4 |
| 13 | Internuron | 82 | 2.2 | 164 | 2.4 |
| 14 | Pyramidal | 87 | 2.3 | 143 | 2.1 |
| 15 | Pyramidal | 68 | 1.8 | 130 | 1.9 |
| 16 | Pyramidal | 76 | 2.0 | 102 | 1.5 |
| 17 | Internuron | 14 | 0.4 | 20 | 0.3 |

**Table S6.** Input resistance is after optoTrkB activation and fluoxetine treatment in WT and hPV-TrkB CKO mice. (A) Input resistance is unchanged after optoTrkB activation. (B) No effect of fluoxetine treatment on input resistance in WT and hPV-TrkB CKO mice.

| Input resistance | Control | 10-30 min | 30-60 min |
| --- | --- | --- | --- |
| Mean | 237.5 | 233.8 | 272.6 |
| Std. Deviation | 63.99 | 58.7 | 77.3 |
| Std. Error of Mean | 26.12 | 22.19 | 31.56 |

| Input resistance | WT water | WT Flx | hPV-TrkB<br>CKO water | hPV-TrkB<br>CKO Flx |
| --- | --- | --- | --- | --- |
| Mean | 171.2 | 251.2 | 189.4 | 204.3 |
| Std. Deviation | 46.84 | 69.45 | 55.23 | 71.69 |
| Std. Error of Mean | 15.61 | 24.55 | 20.87 | 20.69 |

**Supplemental Table 2. Markers in all clusters.**

**Supplemental Table 3. DE genes after optoTrkB activation in all clusters**

DE genes were identified by Chipster/Seurat version 3. Gene ontology (GO) in each gene was annotated by DAVID program

**Supplemental Table 4. Marker genes in clusters of Parvalbumin interneurons**

**Supplemental Table 5. DE genes after optoTrkB activation in clusters of Parvalbumin interneurons.**

DE genes were identified by Chipster/Seurat version 3. GO in each gene was annotated by DAVID program

